# A TLR7/9-IFNα-LDHB axis drives vital NET release and compromises antibacterial defense in lupus

**DOI:** 10.1101/2025.10.07.680928

**Authors:** Eden G. TenBarge, Ashley D. Wise, Morgan L. Hetzel, Helene A. Hoover, Helia Esfandiari, Bailey E. Holder, Eva Belevska, Ellie C. Mennen, Sarah R. McDaniel, Nicole M. Vaccaro, Courtney C. Lucca, Jonathan M. Williams, Jennifer Ferris, Timothy E. Sparer, Leslie J. Crofford, Jeffry D. Bieber, Andrew J. Monteith

## Abstract

Patients with systemic lupus erythematosus (SLE) are susceptible to bacterial infections, but the underlying dysfunction remains unclear. We found that *Staphylococcus aureus* triggers mitochondria-dependent suicidal NETosis via lactate sensing in healthy neutrophils, but this response is defective in SLE. Herein, we show that chronic Toll-like receptor (TLR) 7/9 signaling represses mitochondrial lactate dehydrogenase B (LDHB), thereby impairing lactate sensing and downstream suicidal NETosis. Instead, SLE neutrophils default to vital NET release; a less bactericidal, type I interferon (IFN)-driven process amplified by staphylococcal pore-forming toxins and sustained by elevated systemic IFNα levels observed in SLE. Combined treatment with hydroxychloroquine (HCQ) and interferon-alpha/beta receptor (IFNAR) blockade restores LDHB expression, NET homeostasis, and bacterial clearance in lupus-prone mice. Neutrophils from SLE patients exhibit similar defects, which are reversed by HCQ and the IFNAR-blocking antibody anifrolumab. These findings identify a clinically actionable immunometabolic checkpoint linking chronic autoimmune signaling to defective antibacterial defense in SLE.

## Introduction

Neutrophils are critical for controlling microbial pathogens such as *Staphylococcus aureus.* However, their inflammatory responses must be tightly regulated to balance effective pathogen clearance with host tissue protection. A hallmark of this balance is the formation of neutrophil extracellular traps (NETs), which trap and kill *S. aureus*^1–4^, prevent bacterial dissemination^1,5^, and enhance macrophage activation^6–8^. However, NETs can also drive pathology by exposing nuclear antigens and pro-inflammatory mediators, contributing to autoimmune responses.

In systemic lupus erythematosus (SLE), excessive NET deposition is widely reported and thought to perpetuate disease by promoting type I interferon (IFN) production^9–11^, autoantibody generation^12,13^, endothelial damage^14–16^, thrombosis^17,18^, and cardiovascular disease^19,20^. Impaired NET clearance due to DNase deficiencies^21–23^ or defective lysosomal degradation by macrophages^24–26^ exacerbates these effects, prolonging local inflammation. Human studies similarly associate NET abundance with disease activity in SLE^27–29^, suggesting that dysregulated NET formation and clearance are central to SLE pathology.

NETs are composed of decondensed chromatin decorated with antimicrobial proteins. Functionally, two mechanistically distinct forms exist: suicidal NETosis, in which NETs are released through lytic cell death^2^, and vital NET release, where chromatin is extruded without compromising membrane integrity^3^. Suicidal NETosis has historically been attributed to NADPH oxidase-derived reactive oxygen species (ROS)^30,31^, but recent studies highlighted a role for mitochondrial ROS as an essential trigger, particularly during responses to bacteria^7,32–34^ and in the autoimmune setting of SLE^10,35^. In contrast, vital NET formation is ROS-independent and can be triggered by microbial toxins^3^ or IFNα^36^, although its regulation remains poorly understood. Despite the differences in regulation, both suicidal and vital NET release require the citrullination of histones by protein arginine deiminase 4 (PAD4) to promote DNA decondensation^37^.

Despite their heightened tendency to release NETs, neutrophils from SLE patients fail to undergo mitochondrial ROS-dependent suicidal NETosis in response to *S. aureus*^34,38^. Neutrophils are metabolically unique in that they rely primarily on glycolysis for energy^39–43^, yet they retain functional mitochondria^44,45^. We previously demonstrated that mitochondrial sensing of phagosomal lactate, a byproduct of *S. aureus* fermentation, acts as a critical danger signal that triggers mitochondrial ROS production and initiates suicidal NETosis^34^. In SLE, however, neutrophils exhibit altered mitochondrial homeostasis that impairs this pathway, defaulting instead to chronic vital NET release^34,38^. This paradox suggests that excessive NET formation in SLE may arise from dysregulated mechanisms that favor non-lytic, and potentially less effective, NET programs. This dysregulation may underlie the increased risk of severe bacterial infections in individuals with SLE^46–49^. However, the signaling pathways linking chronic inflammation in SLE to altered neutrophil NET programs remain incompletely understood.

Herein, we identify a critical immunometabolic mechanism by which Toll-like receptor (TLR) 7 and TLR9 signaling represses mitochondrial lactate dehydrogenase B (LDHB) expression in neutrophils, thereby preventing lactate sensing and impairing suicidal NETosis in response to *S. aureus*. Instead, these neutrophils default to vital NET release, a less bactericidal response that is sustained by pore-forming toxins from *S. aureus* and IFNα-driven autocrine signaling. This aberrant NET program is reversed by hydroxychloroquine (HCQ) and interferon-alpha/beta receptor (IFNAR) treatment in lupus-prone mice, restoring LDHB expression, mitochondrial ROS production, and bacterial clearance. Importantly, neutrophils from SLE patients exhibit similar defects in LDHB expression and NET regulation, which are largely corrected by HCQ and IFNAR1-blocking antibody, anifrolumab. These findings reveal a clinically actionable immunometabolic axis linking chronic innate immune activation to defective neutrophil antibacterial responses in SLE.

## Results

### TLR7/9 signaling diminishes LDHB expression and prevents suicidal NETosis in response to S. aureus

We previously demonstrated that reverse conversion of staphylococcal lactate into pyruvate in the mitochondria is essential for triggering mitochondrial ROS and initiating suicidal NETosis^34^. Mammalian cells express two distinct isoforms of LDH: LDHA, which catalyzes the conversion of pyruvate to lactate during glycolysis, and LDHB, which preferentially converts lactate back into pyruvate and associates with mitochondria^50–52^. In line with this, we previously found that mitochondria from SLE neutrophils exhibit reduced LDH activity, specifically, a diminished capacity to convert lactate to pyruvate, compared to healthy controls ^34^. Therefore, loss of LDHB in SLE neutrophils may impair sensing of phagosomal lactate from *S. aureus*, which would compromise suicidal NETosis.

To investigate this, we used MRL/MpJ-*Tnfrs6*^lpr^/J (MRL/*lpr*) mice, a well-established murine model of SLE that recapitulates key autoimmune features of the human disease ^53^. Neutrophils were isolated from MRL/*lpr* and wild-type (WT) C57BL/6 mice, and expression of LDHA and LDHB was assessed by flow cytometry. MRL/*lpr* neutrophils expressed significantly lower levels of LDHB compared to WT, whereas LDHA levels remained unchanged (Fig. 1A, S1A). LDHB mRNA levels were comparable between WT and MRL/*lpr* neutrophils (Fig. 1B), suggesting that LDHB is regulated post-transcriptionally. Consistent with this, mitochondrial fractions from MRL/*lpr* neutrophils also showed reduced LDHB protein abundance by immunoblot (Fig. 1C), indicating loss of mitochondrial LDHB We previously reported that WT neutrophils undergo mitochondrial ROS–dependent suicidal NETosis in response to *S. aureus*, whereas MRL/*lpr* undergo suicidal NETosis in response to apoptotic debris, indicating distinct and potentially antagonistic NET programs^34^. To explore upstream drivers of this divergence, we stimulated WT and MRL/*lpr* neutrophils for 2 hours with agonists for TLR2/1, TLR2/6, TLR3, TLR4, TLR7, and TLR9 in the presence or absence of exogenous lactate. Mitochondrial ROS production and suicidal NETosis were quantified by flow cytometry as previously described^7^. WT neutrophils generated robust mitochondrial ROS (Fig. 2A, S1B) and underwent suicidal NETosis (Fig. 2B, S1C) in response to TLR2 and TLR4 agonists (Pam2CSK4, Pam3CSK4, and LPS) when lactate was present, consistent with their role in bacterial sensing. In contrast, MRL/*lpr* neutrophils responded to TLR7 and TLR9 agonists (IMQ and CPG ODN) with lactate-independent suicidal NETosis, suggesting a shift in NETosis programming.

**Figure 1:**
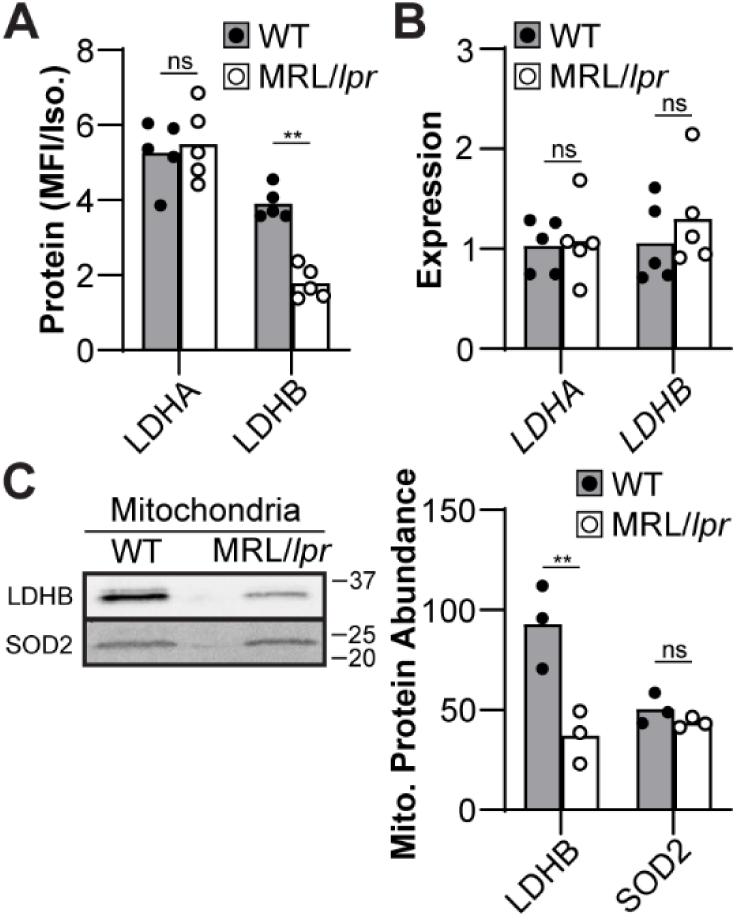
LDHB expression is suppressed in neutrophils from lupus-prone mice. (**A-B**) Neutrophils were isolated from WT or MRL/*lpr* and (*A*) total LDHA or LDHB protein were quantified by flow cytometry relative to an isotype control antibody or (*B*) LDHA or LDHB transcript quantified relative to *ACTB* by quantitative PCR. (**C**) Mitochondria were isolated from WT or MRL/*lpr* neutrophils and the abundance of LDHB or the mitochondrial superoxide dismutase 2 (SOD2) were quantified by immunoblot. Bands were quantified by integrated density. Each point represents neutrophils isolated from a single mouse. (A-B) Two-way ANOVA with Tukey multiple comparisons test or (*C*) paired t test (\*\**p* ≤ 0.01, ns = not significant).

**Figure 2:**
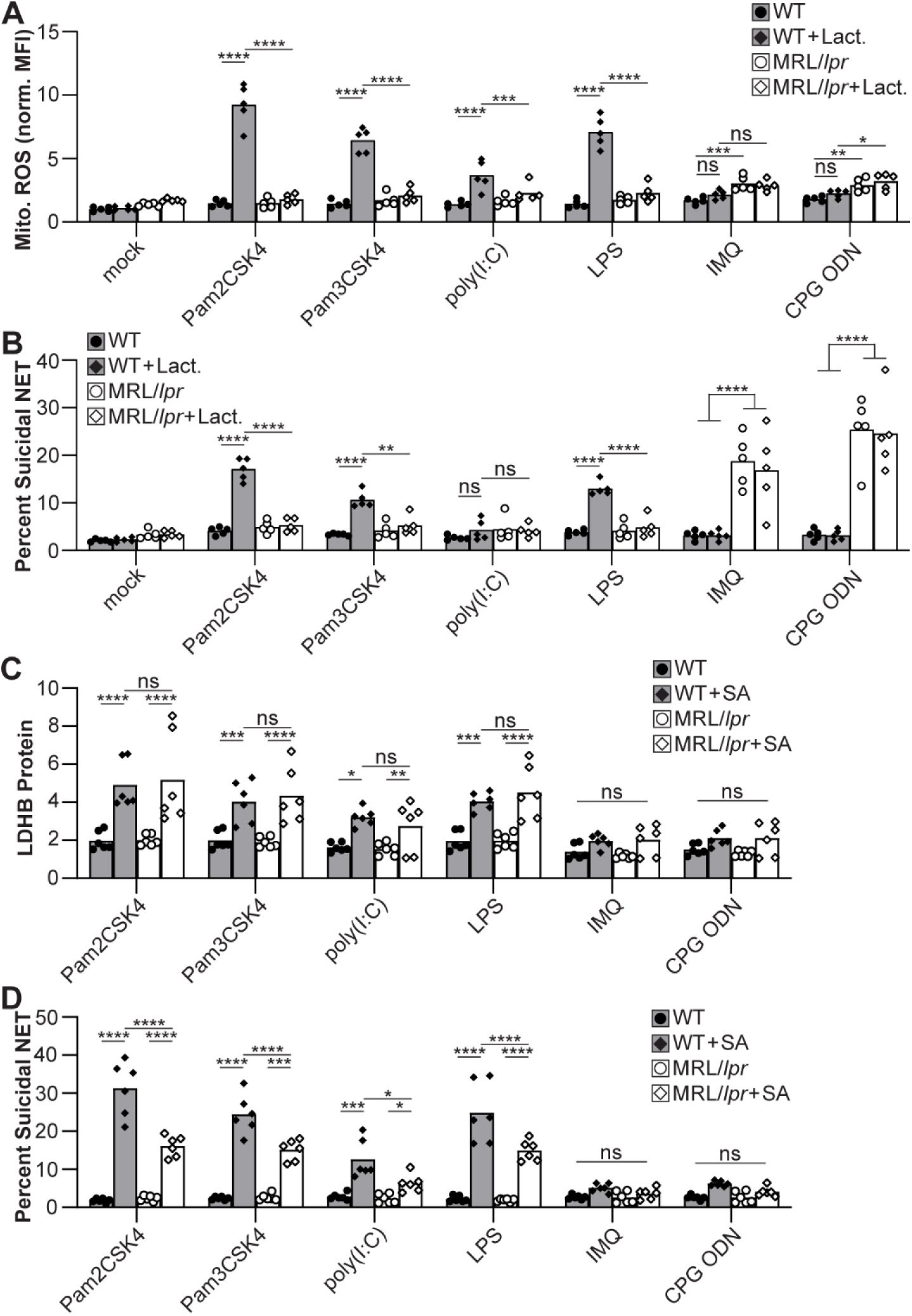
Chronic TLR7/9 signaling suppresses LDHB and impairs lactate-driven suicidal NETosis in lupus-prone neutrophils. (**A**-**B**) WT or MRL/*lpr* neutrophils were treated with differing TLR agonists in the presence or absence of lactate (1 mM) and (*A*) mitochondrial ROS or (*B*) suicidal NETosis were quantified by flow cytometry. (**C**-**D**) Neutrophils were stimulated with TLR agonist (50 ng/mL) and IFNγ (50 ng/mL) and cultured for 16 hours. Neutrophils were cultured with *S. aureus* (SA) for 3 hours and (**C**) total LDHB protein relative to an isotype control antibody or (**D**) suicidal NETosis were quantified by flow cytometry. Each point represents neutrophils isolated from a single mouse. Two-way ANOVA with Tukey multiple comparisons test (\**p* ≤ 0.05, \*\**p* ≤ 0.01, \*\*\**p* ≤ 0.001, \*\*\*\**p* ≤ 0.0001, ns = not significant).

Previous work in LCMV-infected dendritic cells demonstrated that chronic TLR7 activation can suppress LDHB expression^54^. Given that nuclear antigens accumulate in SLE and activate endosomal TLRs^55,56^, including TLR7, we hypothesized that chronic TLR7 stimulation may similarly repress LDHB in neutrophils, impairing their ability to undergo lactate-driven suicidal NETosis. To test this, WT and MRL/*lpr* neutrophils were cultured for 16 hours with IFNγ and TLR agonists (50 ng/mL each), then either mock-treated or stimulated with *S. aureus*. LDHB protein levels, mitochondrial ROS, and suicidal NETosis were subsequently quantified by flow cytometry. In WT neutrophils, overnight stimulation alone did not alter LDHB expression, but pre-treatment with TLR7 or TLR9 agonists followed by *S. aureus* challenge led to significant suppression of LDHB protein (Fig. 2C, S1D) and a corresponding reduction in suicidal NETosis (Fig. 2D, S1E). Conversely, in MRL/*lpr* neutrophils, overnight culture with TLR2 or TLR4 agonists restored LDHB protein levels and partially rescued their capacity to undergo *S. aureus*-induced suicidal NETosis (Fig. 2C–D). These results demonstrate that chronic TLR7 and TLR9 signaling suppresses LDHB expression in neutrophils, impairing their ability to detect bacterial lactate and undergo mitochondrial ROS–dependent suicidal NETosis.

### HCQ restores suicidal NETosis but MRL/lpr neutrophils have impaired antibacterial activity

HCQ, commonly prescribed as Plaquenil, is an acidotropic molecule that binds double- and single-stranded nucleotides and sterically hinders their binding to TLR7/9^57^. Clinically, HCQ is approved for the treatment of SLE and reduces disease flares by dampening type I interferon responses. We previously demonstrated that apoptotic debris triggers aberrant suicidal NETosis in SLE neutrophils^34^. Treating MRL/lpr neutrophils with HCQ prevented this spontaneous NET release in response to apoptotic stimuli (Fig. 3A).

**Figure 3:**
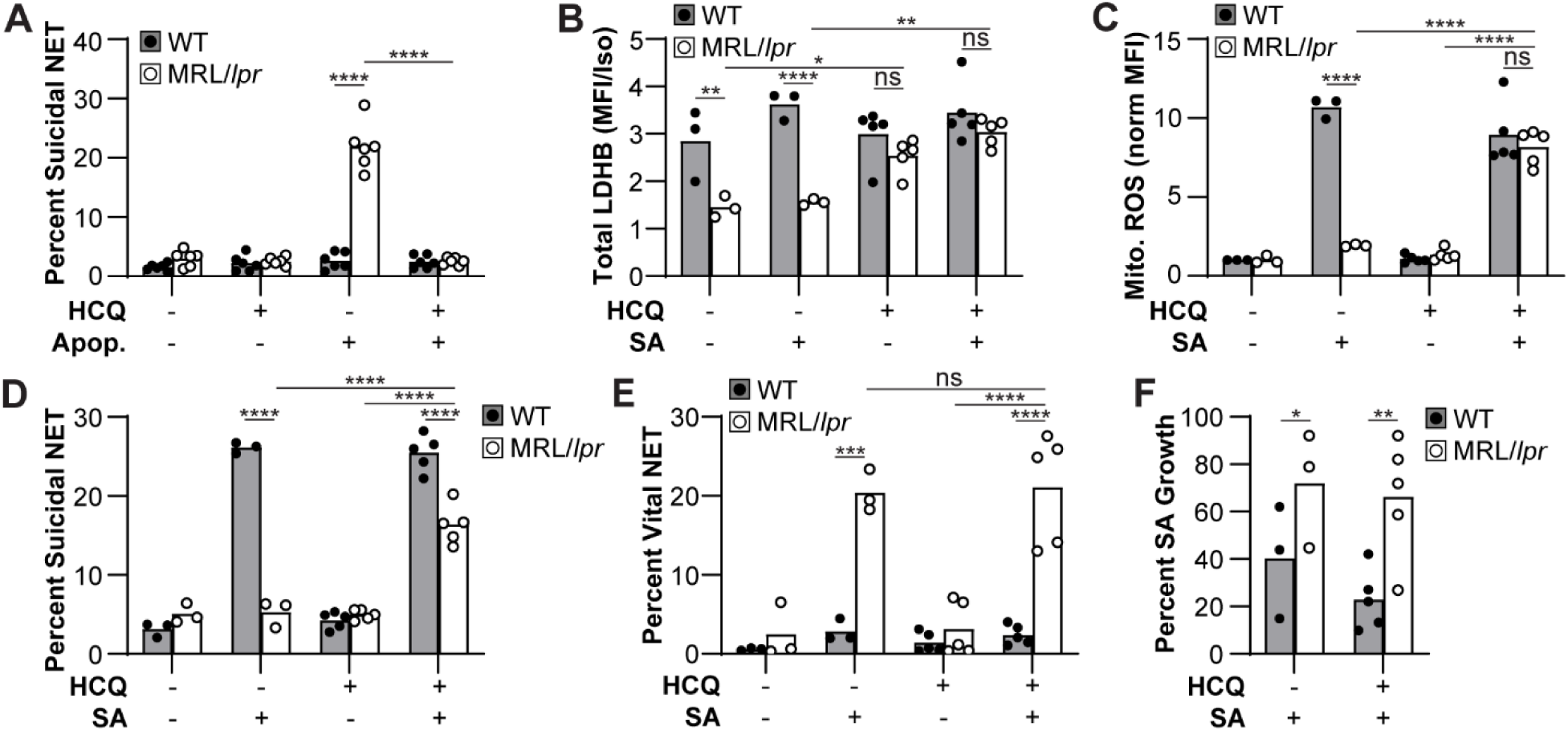
HCQ restores mitochondrial ROS and suicidal NETosis but fails to rescue bactericidal activity in MRL/*lpr* neutrophils. (**A**) WT or MRL/*lpr* neutrophils were treated with HCQ (30 µM) and subsequently stimulated with apoptotic debris. After 3 hours, suicidal NETosis was quantified by flow cytometry. (**B**-**F**) WT or MRL/*lpr* mice were mock treated or treated with HCQ (40 mg/kg) daily for 2 days. Neutrophils were isolated from the bone marrow, cultured with *S. aureus* (SA) for 3 hours, and (*B*) total LDHB protein relative to an isotype control, (*C*) mitochondrial ROS, neutrophils undergoing (*D*) suicidal NETosis or (*E*) vital NET release were quantified by flow cytometry. (*F*) Neutrophils were cultured with *S. aureus* for 4 hours, and bacterial killing relative to *S. aureus* grown in the absence of neutrophils was quantified by dilution spot platting. Each point represents neutrophils isolated from a single mouse. Two-way ANOVA with Tukey multiple comparisons test (\**p* ≤ 0.05, \*\**p* ≤ 0.01, \*\*\**p* ≤ 0.001, \*\*\*\**p* ≤ 0.0001, ns = not significant).

To determine whether HCQ also restores the bacterially induced suicidal NETosis in response to *S. aureus*, WT and MRL/*lpr* mice were treated with HCQ every 24 hours for two days. Neutrophils were then isolated and stimulated *ex vivo* with *S. aureus*. HCQ treatment in MRL/*lpr* mice restored LDHB expression to levels comparable to WT (Fig. 3B, S2A), increased mitochondrial ROS generation in response to *S. aureus* (Fig. 3C, S2B), and re-established the capacity to undergo suicidal NETosis (Fig. 3D, S2C-D).

Despite this restoration of mitochondrial programming, a large subset of MRL/*lpr* neutrophils continued to undergo constitutive vital NET release (Fig. 3E, S2E), a response previously shown to be less effective at bacterial killing^38^. Consistent with this, MRL/*lpr* neutrophils from HCQ-treated mice remained deficient in controlling *S. aureus* infection *ex vivo*, exhibiting impaired bacterial clearance relative to WT controls (Fig. 3F). These findings demonstrate that HCQ treatment can partially rescue suicidal NETosis by restoring LDHB expression and mitochondrial ROS production. However, MRL/*lpr* neutrophils still exhibit sustained vital NET release and remain functionally compromised in their antibacterial response. These data suggest that restoring mitochondrial signaling is necessary but not sufficient for effective bacterial clearance in SLE, potentially due to a persistent bias toward vital NET formation.

### Vital NET release is ineffective at killing S. aureus and interferes with the suicidal NETosis

Although HCQ treatment restores mitochondrial ROS and suicidal NETosis in MRL/*lpr* neutrophils, the continued presence of constitutive vital NET release may compromise antibacterial function. We hypothesized that vital NET release is intrinsically less bactericidal than suicidal NETosis and may actively suppress the latter, either by depleting cellular resources or enforcing a distinct regulatory program.

To experimentally model how one NET program may interfere with another, we established an *in vitro* system to sequentially activate vital and suicidal NET pathways. Neutrophils were pretreated with A23187, a calcium ionophore that triggers vital NET release ^58^, followed by stimulation with phorbol myristate acetate (PMA) or *S. aureus*, both of which induce suicidal NETosis. A23187 pretreatment led to robust vital NET release (Fig. 4A-B) but significantly reduced the fraction of neutrophils undergoing subsequent suicidal NETosis in response to PMA or *S. aureus* (Fig. 4C-D). Conversely, pretreatment with PMA induced suicidal NETosis (Fig. 4C- D) and suppressed subsequent vital NET release in response to A23187 and *S. aureus* (Fig. 4A- B). These findings suggest that vital and suicidal NET programs are mutually antagonistic, consistent with their distinct upstream signaling pathways^58^.

**Figure 4:**
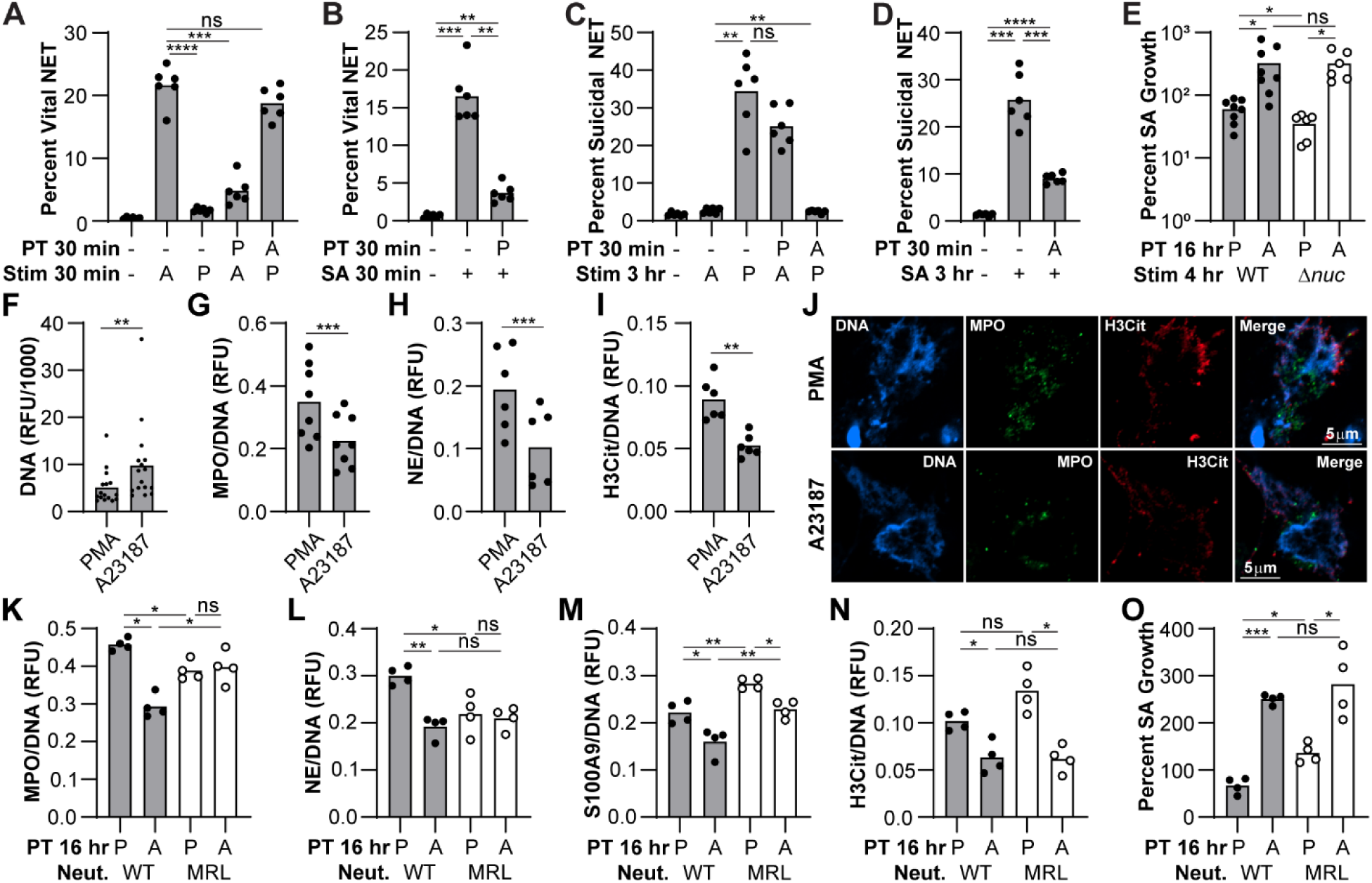
Vital and suicidal NET programs are functionally antagonistic and differ in antibacterial potency and composition. (**A**-**D**) WT Neutrophils were pre-treated (PT) with PMA (P; 5 µM) or A23187 (A; 50 nM) and subsequently stimulated with PMA, A23187, or *S. aureus* (SA). (*A-B*) Vital NET release after 30 minutes or (*C-D*) suicidal NETosis after 3 hours were quantified by flow cytometry. (**E**-**O**) WT and MRL/*lpr* neutrophils were treated with PMA or A23187 for 16 hours and the resulting NETs subsequently (*E, O*) cultured with WT or Δ*nuc S. aureus* for 4 hours. Bacterial killing was quantified relative to *S. aureus* cultured in the absence of NETs by dilution spot plating. (*F*-*N*) Alternatively, NETs were stained to quantify (*F*) DNA, (*G, K*) MPO, (*H, L*) NE, (*M*) S100A9, or (*I, N*) H3Cit using a fluorescent plate reader or (*J*) confocal microscopy. Each point represents (*A*-*D*) neutrophils isolated from a single mouse or (*E*-*I, K-O*) NETs from neutrophils isolated from a single mouse. (*A*-*E, K-O*) One-way ANOVA with Tukey multiple comparisons test or (*F*-*I*) paired t test (\**p* ≤ 0.05, \*\**p* ≤ 0.01, \*\*\**p* ≤ 0.001, \*\*\*\**p* ≤ 0.0001, ns = not significant).

To assess whether these NET programs differ in antibacterial potency, neutrophils were stimulated with A23187 or PMA and cultured for 16 hours to allow NET formation. A23187- induced NETs exhibited significantly reduced capacity to kill *S. aureus* compared to PMA-induced suicidal NETs (Fig. 4E). This difference in bactericidal activity was not due to preferential degradation of suicidal NETs by staphylococcal nucleases, as a nuclease-deficient staphylococcal strain (Δ*nuc*) remained equally resistant to A23187-induced NETs. Despite their impaired antibacterial function, A23187 stimulation led to more abundant NET deposition than PMA (Fig. 4F). However, these vital NETs contained less antimicrobial proteins such as myeloperoxidase (MPO) and elastase (NE), as well as reduced citrullinated histones (H3Cit; Fig. 4G-J). Together, these results demonstrate that vital NETs are both less bactericidal and capable of antagonizing suicidal NETosis.

To further dissect how NET subtype differences contribute to antibacterial efficacy, we compared NET composition and function between WT and MRL/*lpr* neutrophils following stimulation with PMA or A23187. Both genotypes released similar amounts of NETs (Fig. S3A), but their composition diverged. Suicidal NETs from MRL/*lpr* neutrophils contained lower levels of MPO and NE but elevated calprotectin (S100A9) compared to WT (Fig. 4K–M), while vital NETs exhibited modest increases in MPO and calprotectin with similar NE levels. Citrullinated histone content was comparable in vital and suicidal NETs between genotypes (Fig. 4N). Suicidal NETs from MRL/*lpr* neutrophils were less bactericidal than WT (Fig. 4O), consistent with reduced MPO and NE content (Fig. K-L). However, vital NETs from MRL/*lpr* neutrophils remained equally ineffective (Fig. 4O) despite higher MPO and calprotectin levels (Fig. 4K, M). These findings suggest that lupus neutrophils generate less effective suicidal NETs and rely disproportionately on vital NET release, an imbalance that may contribute to impaired bacterial clearance despite excessive NET formation.

### Staphylococcal pore forming toxins cause calcium influx and vital NET release

While the molecular regulation of suicidal NETosis is well characterized, the signaling pathways that drive vital NET release remain poorly defined. Prior work has shown that the pore- forming toxin Panton-Valentine leukocidin (PVL), secreted by *S. aureus*, can trigger vital NET release ^3^. To expand on this, we cultured neutrophils with *S. aureus* mutants lacking either PVL (Δ*pvl*) or α-hemolysin (Δ*hla*), another pore-forming toxin. Both mutants induced significantly less vital NET release compared to WT *S. aureus* (Fig. 5A), confirming that these toxins contribute to vital NET release in WT neutrophils. In contrast, MRL/*lpr* neutrophils released vital NETs independent of these toxins, consistent with their dysregulated NET program.

**Figure 5:**
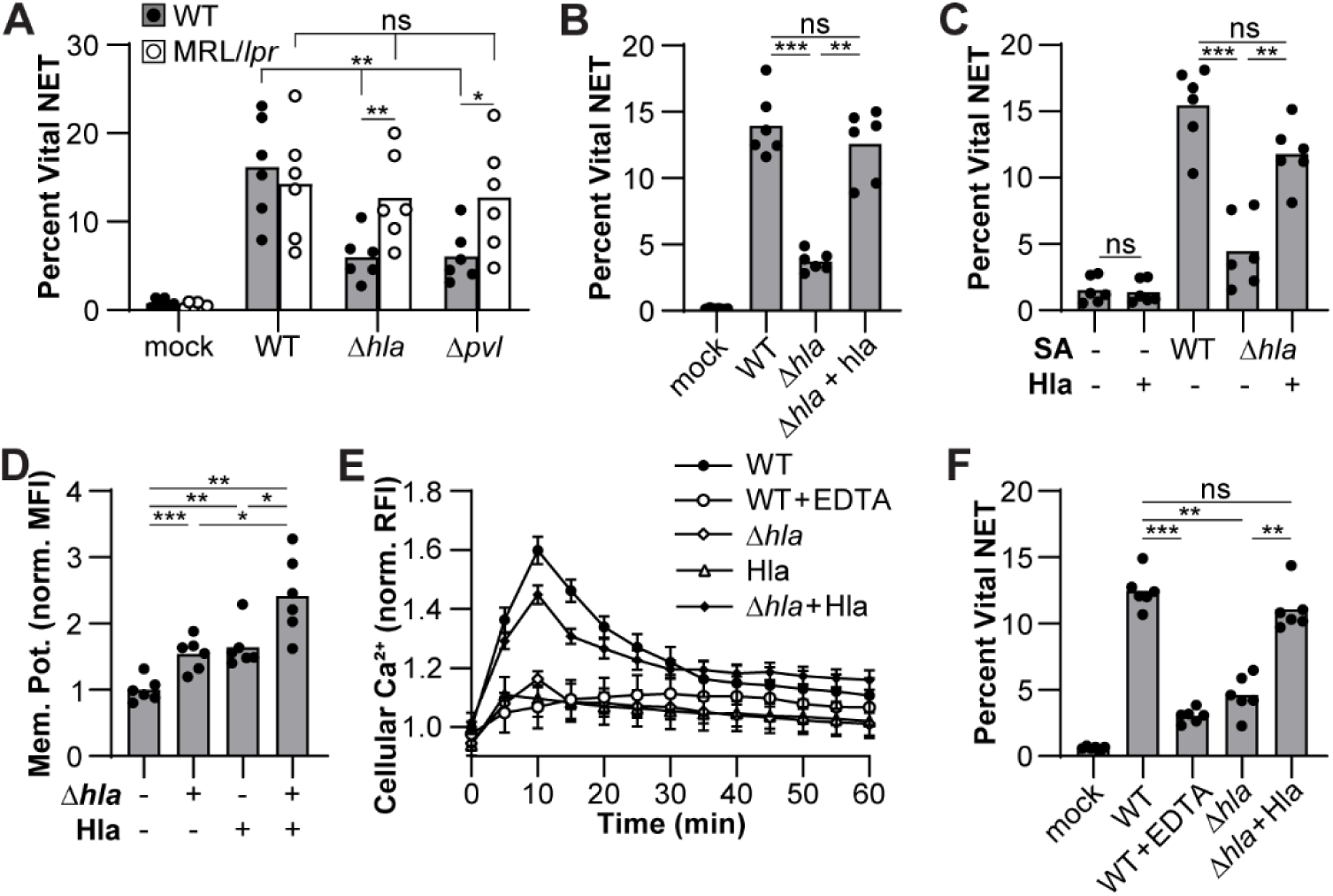
Staphylococcal Hla promotes vital NET release by inducing calcium influx in neutrophils. (**A-C**) WT or MRL/*lpr* neutrophils were cultured with WT, Δ*hla*, Δ*pvl,* or *Δhla* + *hla S. aureus* in the presence or absence of Hla protein and vital NET release was quantified after 30 minutes by flow cytometry. (**D**) WT neutrophils were loaded with DiBAC_4_(3) and stimulated with Δ*hla* and/or Hla protein. The median fluorescence intensity (MFI) was quantified by flow cytometry and normalized to unstimulated neutrophils. (**E**) WT neutrophils were loaded with Fluo-4 and Hoechst 33342 and stimulated with WT or Δ*hla S. aureus* in the presence or absence of EDTA (0.5 mM) or Hla protein (1 µM). Fluo-4 and Hoechst 33342 fluorescence was quantified temporally using a fluorescence plate reader and normalized to unstimulated neutrophils. (**F**) WT neutrophils were cultured with WT or Δ*hla S. aureus* in the presence of EDTA or Hla protein. Vital NET release was quantified by flow cytometry. Each point represents (*A*-*D, F*) neutrophils isolated from a single mouse or (*E*) the mean normalized (Fluo-4/Hoechst 33342) relative fluorescence intensity (RFI) from neutrophils isolated from 5 mouse. (*A*) Two- or (*B*-*D, F*) one-way ANOVA with Tukey multiple comparisons test. (*E*) Error bars = standard error (\**p* ≤ 0.05, \*\**p* ≤ 0.01, \*\*\**p* ≤ 0.001, ns = not significant).

Although PVL promotes vital NET release, we excluded it from further experiments due to its capacity to trigger mitochondrial ROS and suicidal NETosis^33^, which could confound downstream analysis. The Δ*hla* strain did not affect NETosis (Fig. S3B), and thus became the focus of subsequent studies. Expressing Hla in trans (Fig. 5B) or adding exogenous Hla protein (Fig. 5C, S3C) restored vital NET release by WT neutrophils exposed to the Δ*hla* strain. However, Hla protein alone was insufficient to trigger NET release, indicating that it acts in synergy with other bacterial factors. Together, these data demonstrate that *S. aureus* pore-forming toxins, including Hla, are key inducers of vital NET release during infection.

Neutrophils are relatively resistant to Hla-mediated lysis, suggesting that Hla may have immunomodulatory functions beyond cytotoxicity. Differing studies have produced contrasting results where Hla permits calcium influx^59^ or causes depolarization of the cell membrane and prevented calcium influx^60^ leading to enhanced neutrophil inflammatory responses^59,61–63^ but impaired chemotactic migration^60,64^. To test whether Hla alters membrane potential, we loaded neutrophils with the voltage-sensitive dye DiBAC4(3) and stimulated them with the Δ*hla* strain, Hla protein, or both. Each treatment increased DiBAC4(3) fluorescence, indicating membrane depolarization, but the combination of Δ*hla* and Hla protein caused a further increase beyond either treatment alone (Fig. 5D, S3D). This suggests that Hla induces depolarization, likely via sodium influx^60^, and that co-stimulation with *S. aureus* further amplifies depolarization through a distinct ion flux.

Because prior work also implicated Hla in calcium signaling^60^, we tested whether *S. aureus* triggers calcium influx. Neutrophils loaded with the calcium-sensitive dye Fluo-4 showed a rapid and robust increase in intracellular calcium in response to WT *S. aureus*, which was markedly reduced in cells treated with the Δ*hla* strain or calcium chelator EDTA (Fig. 5E). Hla protein alone did not elicit calcium influx, but co-treatment with the Δ*hla* strain restored calcium responses to WT levels, indicating that Hla promotes calcium entry only in the context of bacterial stimulation. Furthermore, vital NET release was diminished when WT neutrophils were stimulated with the Δ*hla* mutant or treated with EDTA (Fig. 5F), underscoring the requirement for both calcium influx and pore-forming toxins. These findings support a model in which Hla-dependent pore formation enables calcium entry into neutrophils, triggering vital NETosis during *S. aureus* infection.

### IFNα sustains vital NET release in MRL/lpr neutrophils responding to S. aureus

IFNα has been shown to induce vital NET release in response to pathogens such as *Mycobacterium tuberculosis*^36^. Given that vital NETs are released within minutes of *S. aureus* stimulation^3^, we hypothesized that IFNα-driven autocrine signaling in this context relies on the rapid secretion of preformed protein rather than de novo transcription or translation. Specifically, we posited that calcium influx triggers degranulation and release of IFNα stores^65–67^. To test this, neutrophils were stimulated with WT or Δ*hla S. aureus*, and surface expression of CD63 and CD35 was quantified by flow cytometry as markers of primary and secretory degranulation, respectively. Primary degranulation in response to *S. aureus* was dependent on the presence of Hla and calcium as the Δ*hla* strain did not trigger and EDTA prevented release of primary granules (Fig. 6A, S3E). In contrast, MRL/*lpr* neutrophils released primary granules in response to *S. aureus* independent of Hla or calcium availability. Secretory degranulation in response to *S. aureus* was dependent solely on the presence of calcium as the Δ*hla* strain triggered release of secretory granules (Fig. 6B). These findings indicate that *S. aureus*-induced degranulation is differentially regulated: primary granule release requires both Hla and calcium, while secretory granule release depends only on calcium.

**Figure 6:**
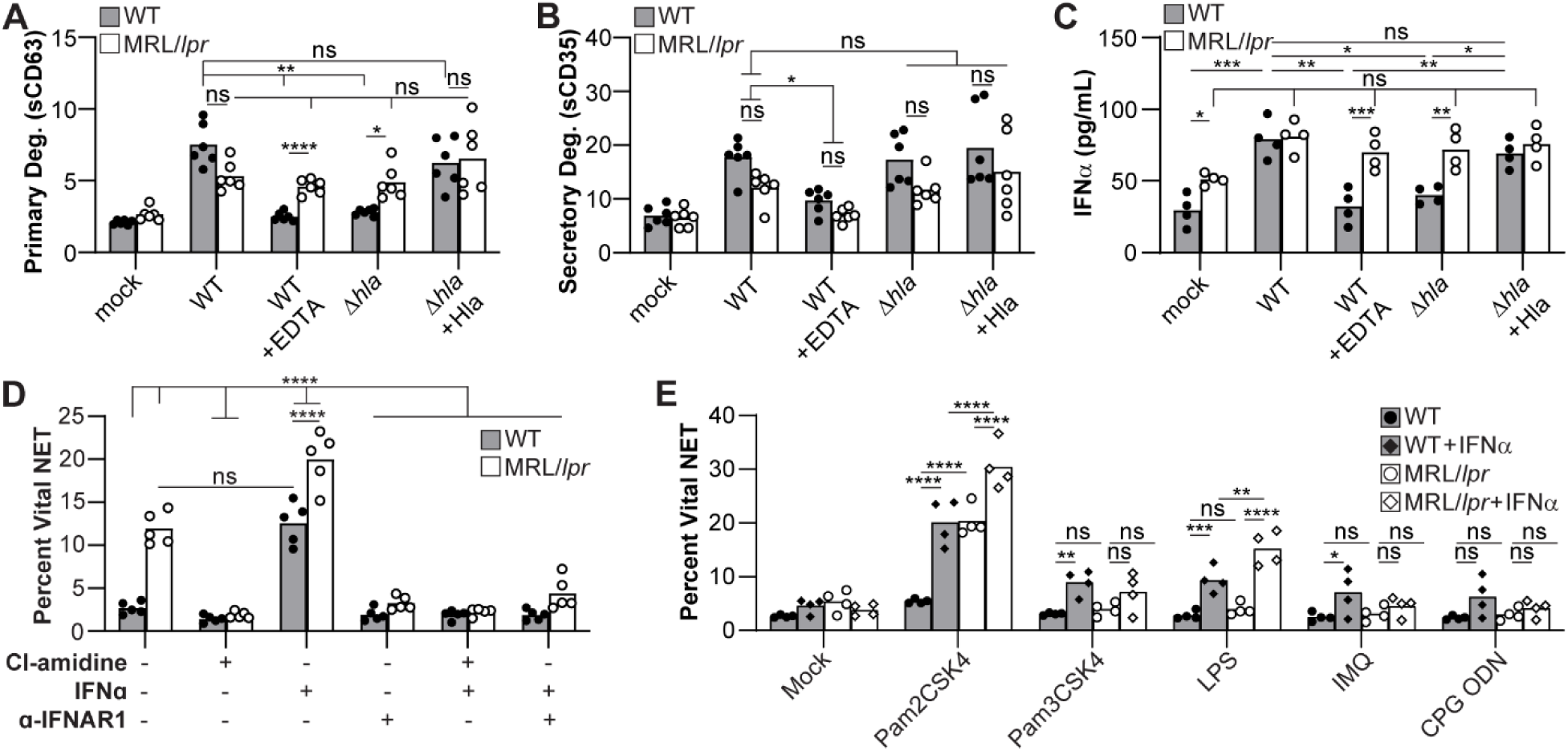
IFNα secretion coincides with calcium-dependent degranulation and is dysregulated in lupus-prone neutrophils. (**A**-**D**) WT or MRL/*lpr* neutrophils were cultured with WT or Δ*hla S. aureus* in the presence or absence of EDTA or Hla protein. After 30 minutes, neutrophils were stained for surface expression of (*A*) CD63 (primary granules) or (*B*) CD35 (secretory granules) and quantified by flow cytometry or (*C*) supernatants isolated to quantify IFNα by ELISA. (**D**-**E**) WT or MRL/*lpr* neutrophils were (*D*) treated with Cl-amidine to broadly inhibit NET release, IFNα, and/or an α-IFNAR1 antibody and cultured with *S. aureus* or (*E*) treated with TLR agonists in the presence or absence of IFNα. After 3 hours, vital NET release was quantified by flow cytometry. Each point represents neutrophils isolated from a single mouse. Two-way ANOVA with Tukey multiple comparisons test (*p ≤ 0.05, **p ≤ 0.01, ***p ≤ 0.001, ****p ≤ 0.0001, ns = not significant).

To determine whether IFNα is released during degranulation, we quantified IFNα secretion by ELISA. WT neutrophils released IFNα in response to WT *S. aureus*, but not to the Δ*hla* strain (Fig. 6C). This defect was rescued by adding Hla protein, confirming that Hla is required for IFNα release. However, calcium chelation with EDTA abolished IFNα secretion, indicating that release is calcium-dependent. This pattern mirrored primary granule degranulation and suggests that IFNα is either stored in primary granules or released concurrently with them. In contrast, MRL/*lpr* neutrophils secreted IFNα regardless of Hla or calcium (Fig. 6C), consistent with their constitutive, dysregulated NET phenotype. Together, these results support a model in which staphylococcal pore-forming toxins initiate IFNα release that drives vital NET release, a pathway that is uncoupled or rendered constitutively active in lupus-prone neutrophils.

We next tested whether IFNα sustains chronic vital NET release. MRL/*lpr* neutrophils released more vital NETs than WT neutrophils after 3 hours of *S. aureus* stimulation (Fig. 6D, S3F), consistent with prior studies^38^. Recombinant IFNα enhanced NET release in both WT and MRL/*lpr* neutrophils, while IFNAR1 blockade or PAD4 inhibition suppressed it. However, IFNα alone was insufficient as TLR co-stimulation was required. In TLR agonist assays, IFNα enhanced NETosis only when paired with a TLR ligand, except in MRL/*lpr* neutrophils responding to TLR2/1, which released NETs even without IFNα (Fig. 6E). These results suggest that TLR2/1 signaling bypasses IFNα dependence in lupus-prone neutrophils and that IFNAR–PAD4 signaling integrates with bacterial sensing to sustain vital NETosis. Overall, our findings support a model in which Hla triggers calcium-dependent IFNα release, which then sustains NET release via IFNAR and PAD4, a program that becomes dysregulated in lupus.

IFNα plays a critical role in driving SLE pathology and similarly, MRL/*lpr* mice have increased serum IFNα (Fig. S4A). Thus, high circulating IFNα may exacerbate vital NET release during infection, impairing the transition to mitochondrial ROS-driven suicidal NETosis. To test this, WT and MRL/*lpr* mice were systemically infected with *S. aureus* by retro-orbital inoculation and treated with either vehicle, HCQ, an anti-IFNAR1 antibody, or both. MRL/*lpr* mice had elevated autoantibody titers and mild glomerulonephritis (proteinuria score ≤ 2) at the time of infection (Fig. S4B-C). Both WT and MRL/*lpr* mice treated with anti-IFNAR1 were highly susceptible to *S. aureus* bacteremia. Dual treatment with HCQ and anti-IFNAR1 in MRL/*lpr* mice did not significantly improve survival (Fig. 7A) despite similar weight loss across treatment groups (Fig. S4D). However, bacterial burdens in the heart and liver were significantly decreased with a strong trend observed in the kidney in dual-treated MRL/*lpr* mice relative to vehicle controls (Fig. 7B). The decrease in bacterial burdens coincided with restoration of LDHB expression (Fig. 7C, S4E), suicidal NETosis (Fig. 7D, S4F), and normalization of vital NET release (Fig. 7E, S4G). These results demonstrate that combined HCQ and IFNAR1 blockade restores mitochondrial function and rebalances NET responses in lupus-prone mice, improving bacterial clearance. However, the modest survival benefit despite improved bacterial control suggests that neutrophil dysfunction is only one facet of infection susceptibility in SLE. These findings underscore the multifactorial nature of immune dysregulation in lupus and highlight the need to address broader inflammatory and tissue-damaging pathways in this setting.

**Figure 7:**
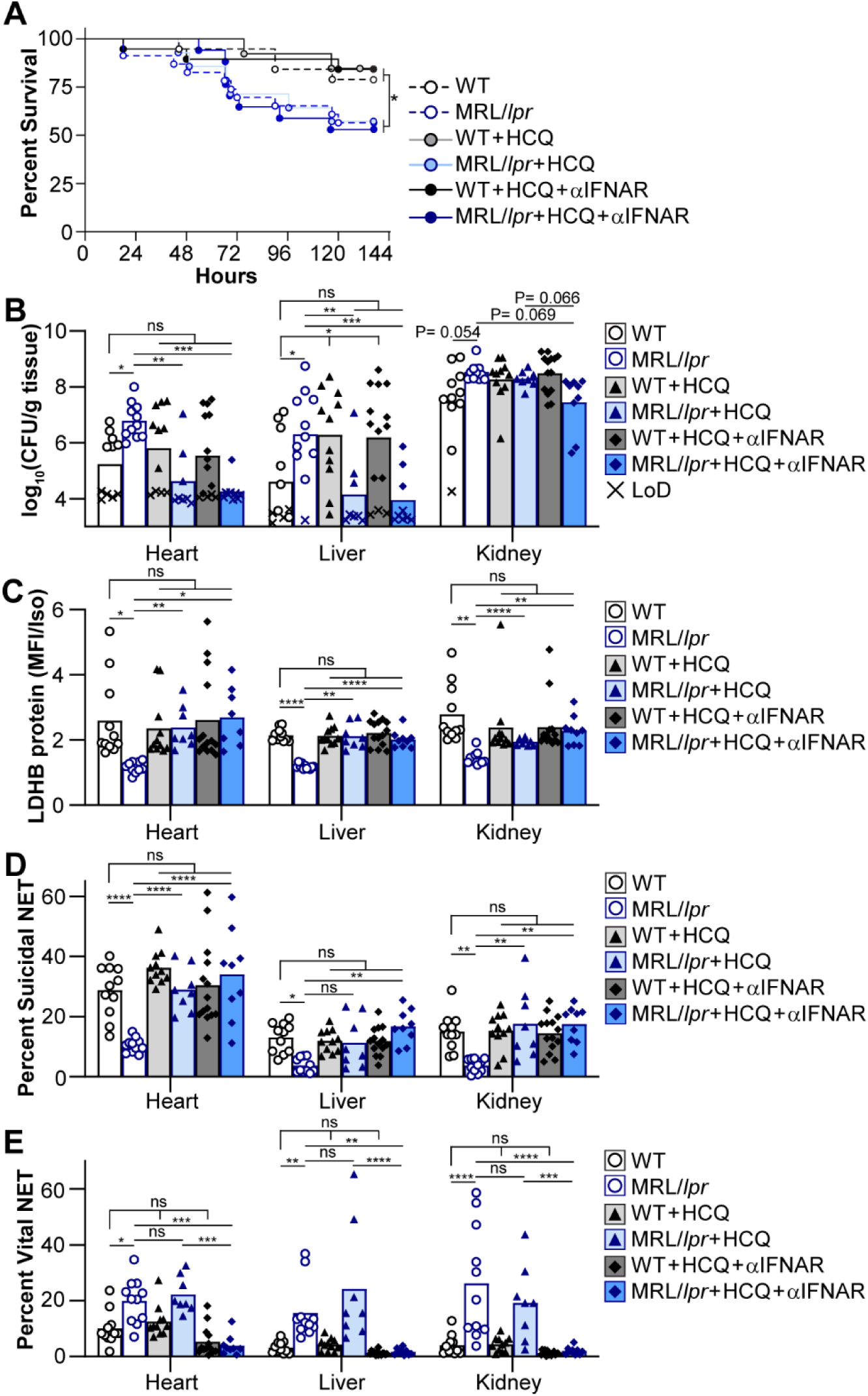
Combined HCQ and IFNAR1 blockade restores NET subtype balance and antibacterial defense in lupus-prone mice. WT or MRL/*lpr* mice were retro-orbitally infected with *S. aureus* and treated with vehicle, HCQ (40 mg/kg/ every other day), and/or anti-IFNAR1 antibody (10 mg/kg/every other day). (**A**) Mouse survival was monitored daily post-infection. At 6 days post-infection, (**B**) bacterial burdens in the heart, liver, and kidney were quantified by dilution spot plating and neutrophils from the tissues analyzed for (**D**) LDHB protein expression relative to isotype control, (**E**) mitochondrial ROS, (**F**) suicidal NETosis, and (**G**) vital NET release by flow cytometry. (*A*) Data represents the mean survival of mice and (*B*-*G*) each point represents a single mouse. (*A*) Log-rank (Mantel–Cox) test and (*B*-*G*) two-way ANOVA with Tukey multiple comparisons test (*p ≤ 0.05, **p ≤ 0.01, ***p ≤ 0.001, ****p ≤ 0.0001, ns = not significant).

### HCQ and Anifrolumab therapies restore regulation of NET release by SLE neutrophils

MRL/*lpr* mice are genetically identical and housed in controlled conditions, whereas SLE in patients arises from diverse genetic and environmental influences. To determine whether the mechanisms identified in lupus-prone mice extend to human disease, we analyzed neutrophils from individuals with SLE (Tables S1-S2). Neutrophils were isolated from healthy controls (HC) and SLE patients, and LDHB protein expression was quantified by flow cytometry. SLE neutrophils from patients receiving higher doses of HCQ, normalized to body weight, expressed significantly more LDHB compared to those on lower doses, though levels remained reduced relative to HC neutrophils (Fig. 8A–B). *Ex vivo* treatment of SLE neutrophils with *S. aureus* and HCQ increased LDHB expression to levels comparable to HC controls. Given this rescue of LDHB, we next assessed mitochondrial ROS production (Fig. 8C–D) and suicidal NETosis (Fig. 8E–F) in response to *S. aureus*. We analyzed responses both in freshly isolated neutrophils and by retrospectively correlating HCQ dosing with prior data^34,38^. SLE neutrophils from patients on higher doses of HCQ produced more mitochondrial ROS and underwent greater suicidal NETosis following *S. aureus* challenge (Fig. 8C, E). In addition, *ex vivo* HCQ treatment restored both responses to levels seen in HC neutrophils (Fig. 8D, F). These findings indicate that HCQ dose-dependently restores LDHB expression and mitochondrial programming, thereby rescuing the capacity for suicidal NETosis in SLE neutrophils.

**Figure 8:**
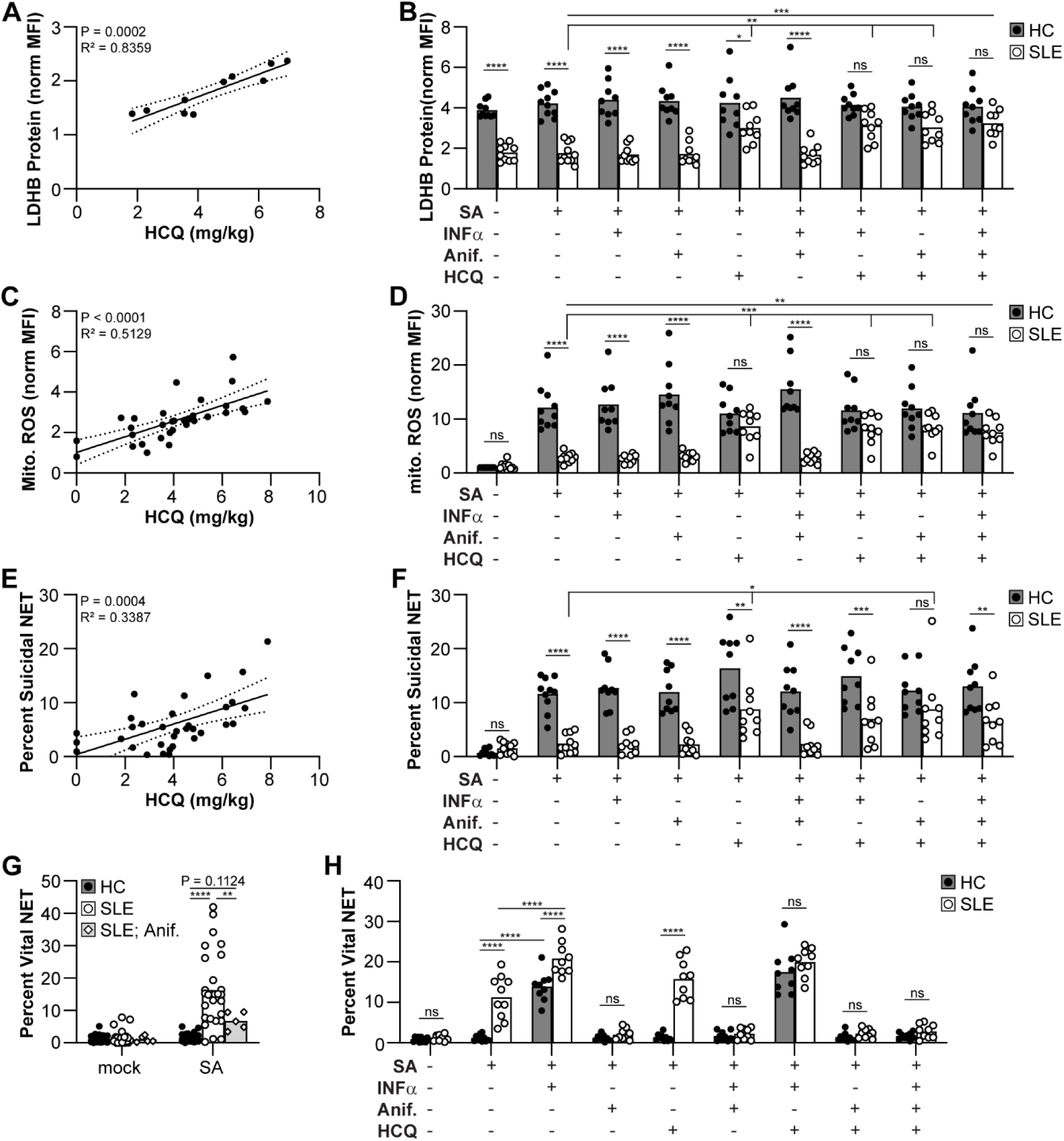
HCQ and anifrolumab therapies restore NET subtype regulation in SLE neutrophils. Neutrophils were isolated from HC or SLE patients. (**A**-**B**) LDHB protein expression relative to isotype control in unstimulated neutrophils, or (**C**-**D**) mitochondrial ROS, (**E**-**F**) suicidal NETosis, (**G**-**H**) vital NET release in response to *S. aureus* were quantified by flow cytometry. (*A, C, E*) LDHB expression, mitochondrial ROS, and suicidal NETosis was graphed relative to current dosage of HCQ. (*G*) Vital NET release were quantified in patients not receiving or receiving anifrolumab (anif.) treatment. (B, D, F, H) Neutrophils were treated with IFNα, HCQ, and/or anifrolumab and stimulated with *S. aureus* for 3 hours. Each point represents neutrophils isolated from a single donor. (*A*, *C*, *E*) Data were quantified by linear regression or (*B, D, F-H*) one-way ANOVA with Tukey multiple comparisons test (*p ≤ 0.05, **p ≤ 0.01, ***p ≤ 0.001, ****p ≤ 0.0001, ns = not significant). (*C, E, G*) Includes data from this study and prior studies ^34,38^.

We next assessed chronic vital NET release in response to *S. aureus*. Neutrophils from SLE patients receiving anifrolumab exhibited significantly less vital NET release compared to untreated patients (Fig. 8G). Furthermore, anifrolumab suppressed vital NET formation to levels observed in HC neutrophils, while exogenous IFNα augmented vital NET release in HC neutrophils to levels resembling SLE (Fig. 8H). Together, these findings demonstrate that the aberrant NET program observed in lupus-prone mice is recapitulated in human SLE neutrophils and can be pharmacologically reversed. HCQ restores mitochondrial function and suicidal NETosis, while anifrolumab suppresses IFNα-driven vital NET release, providing mechanistic insight into how standard-of-care therapies modulate innate immune dysfunction in SLE.

## Discussion

Neutrophils are essential for early antibacterial defense, yet in SLE, their responses become paradoxically maladaptive. Excessive NET formation fuels autoimmunity^27–29^, while susceptibility to infections like *S. aureus* remains high^46–49^. This contradiction suggests that not all NETs are equal in function. Here, we reveal that chronic innate immune activation in lupus reshapes NET output from bactericidal suicidal NETosis to a constitutive, ineffective vital NET program, undermining host defense despite increased NET burden.

We identify a TLR7/9-IFNα-LDHB immunometabolic axis that governs neutrophil NET programming. Similar to LCMV-infection in dendritic cells^54^, chronic TLR7/9 stimulation during SLE represses LDHB, a mitochondrial enzyme required for lactate sensing and ROS production during bacterial infection^34^. This repression severs a key metabolic checkpoint, blocking mitochondrial ROS and suicidal NETosis in response to *S. aureus*. Instead, lupus-prone neutrophils default to vital NET release, a program we show is less bactericidal and actively antagonizes suicidal NETosis, likely through pathway competition and granule depletion.

We further elucidate the upstream regulation of vital NET release. Staphylococcal Hla facilitates calcium influx and degranulation, initiating rapid IFNα secretion and vital NET formation. In healthy neutrophils, this occurs via a tightly regulated process that depends on intact calcium signaling. This is consistent with earlier work demonstrating that *S. aureus* triggers rapid, vesicle-mediated NET release via non-lytic, oxidant-independent pathways, particularly in response to PVL and related pore-forming toxins^3^. More recently, type I interferon was shown to enable neutrophils to extrude NETs while maintaining membrane integrity in TB granulomas^36^. These findings align with our observation that IFNα signaling amplifies vital NET release and suggest that this pathway, while beneficial in some contexts, may become pathologically unrestrained in SLE. In lupus-prone neutrophils, IFNα release becomes decoupled from calcium dependence, enabling constitutive autocrine IFNAR signaling that sustains vital NET formation. Our findings reconcile previously conflicting reports on Hla’s effects on calcium flux and neutrophil activation^59–64^, and position vital NETosis as a context-dependent, IFNα-amplified program that contributes to defective host defense in lupus.

This NET subtype imbalance has functional consequences. Vital NETs lack the antimicrobial protein content and killing capacity of suicidal NETs, yet are produced in greater abundance. In lupus-prone neutrophils, this imbalance not only fails to control *S. aureus* but also suppresses the more effective suicidal program, compounding infection risk. Both vital and suicidal NETs from MRL/lpr neutrophils contained elevated levels of S100A9, consistent with the increased calprotectin expression we previously reported in lupus neutrophils^38^. Although calprotectin contributes to antibacterial defense^68,69^, our findings demonstrate that its enrichment alone is insufficient to compensate for the loss of effective suicidal NETosis. These results align with recent evidence that *S. aureus* PVL triggers vital NET formation, yielding NETs with reduced NE, cathelicidin, and proteinase 3 content and poor bactericidal activity^70^. Instead, the skew toward vital NET release exacerbates functional defects, impairing bacterial clearance despite excessive NET deposition. These findings resolve a longstanding paradox in lupus pathogenesis: NET abundance is not protective when the dominant subtype lacks bactericidal potency.

Importantly, this dysregulated NET program is therapeutically reversible. In lupus-prone mice, HCQ restores LDHB expression, mitochondrial ROS, and suicidal NETosis by blocking endosomal TLR7/9 signaling. IFNAR blockade with anti-IFNAR1 antibody disrupts IFNα-driven vital NET release. Combined therapy rebalances NET subtypes and reduces bacterial burden *in vivo.* In human SLE neutrophils, defects in LDHB and NET programming mirror those in mice, and are corrected by HCQ or anifrolumab, providing mechanistic rationale for these therapies and their benefits to innate immunity.

However, rebalancing NET responses alone was insufficient to improve survival in septic lupus-prone mice, highlighting the multifactorial nature of infection vulnerability in SLE. Factors such as impaired macrophage function^25,26,71–74^, vascular pathology^14–20^, and complement deficiencies^75^ likely contribute to poor outcomes. Moreover, it remains to be seen whether similar NET imbalances impair responses to other pathogens, including viruses and fungi, particularly given the critical role of IFN signaling in antiviral defense.

Our findings underscore several broader implications. First, NET subtype identity is functionally decisive, not merely morphological. Suicidal NETosis is linked to robust antimicrobial content and cell death, whereas vital NETs are non-lytic, less antimicrobial, and suppress more effective NET programs. Second, LDHB emerges as a central metabolic node linking innate immune signaling to mitochondrial fitness and effector function, one that may be dysregulated across multiple cell types and diseases marked by chronic inflammation^76–79^. Finally, NET subtype composition may represent a tractable biomarker of neutrophil health and treatment response in SLE and other inflammatory diseases.

In summary, this study identifies a TLR7/9-IFNα-LDHB axis that rewires neutrophil NET programming in lupus. By defining how chronic inflammation impairs antibacterial immunity through metabolic and signaling reprogramming, we reveal an actionable mechanism that reconciles key paradoxes in SLE pathogenesis and lays the groundwork for targeting innate dysfunction in autoimmune disease.

## Methods

### Reagents

Antibodies specific for Ly6G (127643), CD15 (323050), CD16 (302016), CD63 (143903), CD21/35 (123409), and PE isotype control (400508) were from Biolegend. CD11b (60-0112- U100) was from Tonbo and LDHA (PA523036) and LDHB (MA541126) were from Invitrogen. Streptavidin Alexa Fluor 488 (405235), rabbit Alexa Fluor 488 (405235), and rabbit Alexa Fluor 647 were purchased from Biolegend. MPO (AB90810), H3Cit (AB5103), and Neutrophil Elastase (AB68672) were from Abcam and S100A9 (73425) was from Cell Signaling. Murine Fc Shield (70- 0161-M001) was purchased from Tonbo and Human TruStain FcX (422302) was from Biolegend. Paraformaldehyde (15710) was from Electron Microscopy Sciences. MitoSOX red (M36008), MitoTracker deep red (M22426), SYTOX blue (S34857), DiBAC4(3) (B438), Cell Mask deep red (C10046), and prolong gold (P36930) were purchased from Invitrogen. Hoechst 33342 (62249) and Fluo-4 (F14201) were from Thermo Scientific and Ghost Dye Violet 540 (13-0879-T100) was from Tonbo. Histopaque 1077 (10771-6X100ML), Histopaque 1119 (11191-6X100ML), and Poly-L Lysine (P4707) were purchased from Sigma-Aldrich. DMEM (11-965-118) and FBS (A52567- 01, Lot 2635802RP) were from Gibco, DPBS (16750-102) was from Cytiva, and 10X RBC lysis buffer (420302) was from Biolegend. α-dsDNA (AB27156) and IgG AP (AB98724) were purchased from Abcam and salmon sperm DNA solution (15632011) was from Invitrogen. Laemmli loading buffer (1610747) was from BioRad and the chemiluminescent substrate kit (34580) was from Thermo Scientific. The antibodies SOD2 (D3X8F) and goat anti-rabbit IgG HRP (W401B) were from Cell Signaling and Promega, respectively. The mitochondrial isolation kit (89874) was purchased from Thermo Scientific and the mouse IFNα ProQuantum Immunoassay kit (A46736) was from Invitrogen. The Turbo DNAse kit (AM1907), RNA HS kit (Q32852), and SuperScript III first strand synthesis kit (18080051) were purchased from Invitrogen. The RNeasy mini kit (74104) and Proteinase K (19131) were from Qiagen. Mutanolysin (M9901-1KU) was from Sigma, lysozyme (90082) was from Thermo Fisher, and EDTA (BP2482100) was from Fisher. TaqMan Gene Expression Master Mix (4369514) was purchased from Applied Biosystems. NaLactate (71718), PMA (P8139), A23187 (AC328080050), Cl-Amidine (506282), Hla protein (H9395), and LPS (L4391-10) were purchased from Sigma-Aldrich. Pam2CSK4 (4637), Pam3CSK4 (4633), Poly I:C (4287), and Imiquimod (3700) were from Tocris. Mouse IFNγ (315- 05), and human IFNα (300-02AA) were from PeproTech. CpG ODN (TLRL1826) was purchased from Invivogen. Mouse α-IFNAR (127324) and Mouse IFNα (752806) were from Biolegend and human α-IFNAR (A2460) was from Selleck.

### Bacteria

All *S. aureus* experiments used the bacterial strain USA300 (LAC). The Δ*nuc* mutant ^1^ was from the Horswill lab. The ΔPVL strain was from the Nebraska Transposon Mutant Library ^80^ and the Δ*hla* mutant ^81^ was from Megan Kiedrowski. Bacteria were maintained as a -80°C glycerol stock, and *S. aureus* was streaked onto tryptic soy agar (TSA; 2% agar) plates to grow overnight at 37°C. Single colonies were transferred to tryptic soy broth (TSB) and liquid cultures were grown overnight at 37°C with 250 rpm of shaking. For all in vitro experiments, overnight cultures were diluted 1:50 into non–heat-inactivated FBS before coculture with neutrophils.

### Mice

C57BL/6J and MRL/MpJ-*Fas^lpr^*/J mice were purchased from the Jackson Laboratory (JAX mice stock no. 000664 and no. 000485). A C57BL/6J breeding colony was subsequently maintained in an accredited animal facility. Mice were housed two to five mice per cage with an independently ventilated air supply and food and water were provided ab libitum. Cages were changed on a weekly basis. For hydroxychloroquine (HCQ) treatments, mice were injected intraperitoneally at a concentration of 40 mg/kg every 24 hours for 2 days prior to neutrophil isolation. Mock treatment was the diluent (DPBS). All experiments were conducted in accordance with protocols approved by the University of Tennessee Institutional Animal Care and Use Committee.

### Human neutrophil isolation

Neutrophils were isolated from the peripheral blood of donors using density centrifugation where neutrophils were isolated at the interface between layers of Histopaque 1119 and Histopaque 1077. Serum was pulled off the top of the blood prior to density centrifugation each experiment for opsonization of bacteria. Blood was diluted with DPBS then layered on top of 2-3 Histopaque stacks. Isolated neutrophils were washed with DMEM and then incubated on ice for 45 minutes in DMEM with 10% FBS (D10 media) before lysing red blood cells with 1-2mL 1X RBC lysis buffer. The isolated neutrophils rested 1 hr at 37°C, 5% CO_2_ before experiments. The isolated cells were 90-95% neutrophils (CD15^+^CD16^+^), as verified by flow cytometry.

### Murine neutrophil isolation

Single-cell suspensions of bone marrow were extracted from the tibias and femurs of female mice aged 6-10 weeks and layered on top of a Histopaque stack. Following density centrifugation, the neutrophils were isolated at the interface of Histopaque 1077 and Histopaque 1119. The cells were washed with DMEM and resuspended in D10 media. The neutrophils were rested on ice for 1 hr, plated, and allowed to rest for 1 hr at 37°C, 5% CO_2_ before experiments. Alternatively, neutrophils were cultured with IFNγ and TLR agonists (50 ng/mL) for 16 hours at 37°C, 5% CO_2_ before conducting experiments. The isolated cells were 85-95% neutrophils (CD11b^+^Ly6G^+^), as verified by flow cytometry.

### Apoptotic Debris

Following euthanasia, the thymus was removed from a mouse less than 15 weeks of age, grinded through a 70 µm cell strainer and rinsed with DPBS. Cells were irradiated (6 gray), placed in a 6 cm petri dish, and incubated at 37°C, 5% CO_2_ overnight. The next day the sample was transferred to a 15 mL conical tube and pelleted at 350 x g for 5 min. Supernatant containing apoptotic debris was transferred to a fresh tube and added to serum to stimulate neutrophils.

### Murine bacteremia models

*S. aureus* was streaked from frozen stocks on TSA. Overnight cultures of *S. aureus* were grown in TSB and then subcultured 1:100 in 5 mL of TSB. *S. aureus* was grown for 3 hours, washed, and resuspended in ice-cold DPBS. Bacteria was resuspended at 2×10^8^ CFU/mL for C57BL/6 or 4×10^8^ CFU/mL for MRL/*lpr* to account for the differences in size between the two mice. All infections used 10- to 12-week-old female mice, which were anesthetized using inhalation of isoflurane followed by inoculation of the retro-orbital sinus with 100 µL of bacterial slurry or mock treated with 100 µL DPBS. A lubricant eye ointment was applied after each injection and mice were monitored until they regained consciousness. Mice were monitored daily before humane euthanasia by inhalation of CO_2_.

For anti-IFNAR (anifrolumab) and hydroxychloroquine (HCQ) treatments, mice were injected intraperitoneally every other day beginning the day after infection. Anti-IFNAR was administered at a concentration of 10 mg/kg and HCQ at 40 mg/kg. Mock treatments were the respective diluent.

One day prior to infection, murine levels of proteinuria and serum autoantibody were quantified. Urine protein was assessed according to Uristix 4 (Siemens) instruction. Blood was collected by tail nick and serum levels of IgG reactive to dsDNA was quantified by ELISA.

### Organ Harvest

Following euthanasia, the heart, liver, and kidney were removed, weighed and placed into tubes containing DPBS on ice. Blood was collected from cardiac sticks for IFNα ELISA. Single cell suspensions were created by grinding organs through 70 µm cell strainers and rinsing in DPBS. Aliquots of 200 µL of the single cell suspension were reserved for serial dilution spot plating before centrifugation to determine bacterial load. Cells were pelleted and the pellets resuspended with 1X RBC lysis buffer per manufacturer’s instructions. Cells were strained through a 70 µm cell strainer, pelleted by centrifugation, washed in DPBS, and resuspended in FACs media (DPBS with 2% heat inactivated FBS and 0.02% NaAz) and plated for flow cytometry.

### Bacterial Burdens from Infections

Samples were aliquoted into the top row of a 96 well plate and then serially diluted in DPBS using 10-fold increments down the plate. 5 µL of each dilution was then spotted onto TSA plates and incubated for 12-16 hours in a static 37°C incubator. Plates were removed from the incubator and CFU counted.

### Flow Cytometry

An Attune NXT flow cytometer was used to run all flow cytometry assays. All data was analyzed using FlowJo software. For all flow cytometry data, cells were gated side scatter height (SSC-H) by side scatter area (SSC-A) followed by forward scatter height (FSC-H) by forward scatter area (FSC-A) to remove doublet populations. The singlet population was gated SSC-H by FSC-H to isolate the granulocyte populations. The resulting cell population was then assessed for assay-specific fluorescent markers as described below.

#### Exogenous Treatments

Neutrophils were pre-treated with HCQ (30 pM) 30 minutes prior to stimulation and EDTA (0.5mM) 5 minutes prior to stimulation. Neutrophils were stimulated with bacteria at an MOI = 10. Cl-amidine (34 µg/mL), IFNα (200 units/well), α-IFNAR (10 µg/mL), Pam2CSK4 (1 µg/mL), Pam3CSK4 (1 µg/mL), Poly I:C (20 µg/mL), LPS (1 µg/mL), imiquimod (2 µg/mL), CpG ODN (45 nM), sodium lactate (1 mM), and/or Hla protein (1 µg/mL) were added concurrently with bacteria. A23187 (5 µM) and PMA (50 nM) were added to trigger vital NET release or suicidal NETosis. Alternatively, A23187 or PMA were added 30 minutes prior subsequent simulations or bacteria.

#### Mitochondrial ROS Flow

Following plating, neutrophils were stained with mitoTracker (500 nM) deep red and mitoSOX red (5 µM) dyes for 20 min, washed, and fresh D10 media added to the neutrophils. Neutrophils were stimulated for 3 hours. Neutrophils were stained with Ghost Dye™ Violet 540 to monitor membrane integrity 20 min prior to fixation. Neutrophils were fixed with room temperature 4% PFA and placed on ice for 20 min. Neutrophils were pelleted and blocked in FACS media (mouse = Fc Shield; human = Human TruStain FcX) on ice for 20 min. Neutrophils were pelleted, and resuspended in FACS media containing fluorescent antibodies (mouse = α-CD11b, α-Ly6G; human = α-CD16, α-CD15) for 20 min on ice. Neutrophils were washed and resuspended in FACS media for analysis. Cells were gated for live cells (Ghost^-^) and then neutrophils (CD11b^+^Ly6G^+^ or CD15^+^CD16^+^). mitoSOX and mitoTracker mean fluorescence intensity (MFI) were normalized to untreated or mock WT MFI.

#### NETosis Flow

NETosis flow was conducted as previously described ^7^. Neutrophils were stimulated for 30 minutes or 3 hours and Ghost Dye™ Violet 540 and SYTOX blue (500 nM) were added to the culture 20 min prior to fixation to monitor membrane integrity and extracellular DNA, respectively. Neutrophils were fixed with room temperature, 4% PFA for 20 min on ice and subsequently blocked in FACS media (mouse = Fc Shield, human = Human TruStain FcX) on ice for 20 min. Neutrophils were pelleted and resuspended in FACS media containing fluorescent antibodies (mouse = α-CD11b, α-Ly6G; human = α-CD16, α-CD15) and biotinylated α- myeloperoxidase and rabbit α-H3Citruline for 20 min on ice. Neutrophils were pelleted and resuspended in FACS media containing fluorescently-labeled streptavidin and α-rabbit antibodies on ice for 20 min. Neutrophils were washed and resuspended in FACS media for analysis. Cells were gated for neutrophils (CD11b^+^Ly6G^+^ or CD15^+^CD16^+^) and cells positive for extracellular dsDNA (SYTOX^+^), MPO, and H3Cit were defined as having NET release. Neutrophils releasing NETs that have permeabilized membranes (Ghost^+^) were defined as having undergone suicidal NETosis, while Neutrophils releasing NETs that do not have permeabilized membranes (Ghost^-^) were defined as having undergone vital NET release. The percentage of neutrophils undergoing NETosis was quantified relative to the total number of neutrophils.

#### LDH Protein Flow

To quantify intracellular LDH abundance, neutrophils were stained with Ghost Dye™ Violet 540 20 min prior to fixation, fixed with room temperature 4% PFA for 20 min on ice, blocked in FACS media (mouse = Fc Shield; human = Human TruStain FcX) on ice for 20 min, and stained in FACS media containing fluorescent antibodies (mouse = α-CD11b, α-Ly6G; human = α-CD16, α-CD15) on ice. Neutrophils were then stained with a rabbit α-LDHA or -LDHB antibody or isotype control in Saponin Permeabilization Buffer for 20 min on ice, and stained with a fluorescent α-rabbit antibody in Saponin Permeabilization Buffer for 20 min on ice. Neutrophils were washed and resuspended in FACS media for analysis. LDH MFI was normalized to an isotype control.

#### Degranulation Flow

Neutrophils were stimulated for 30 minutes and stained with Ghost Dye™ Violet 540 20 min prior to fixation, fixed with room temperature 4% PFA for 20 min on ice, blocked in FACS media (Fc Shield) on ice for 20 min, and stained with antibodies recognizing appropriate neutrophil surface markers (murine: CD11b, Ly6G; human: CD15, CD16), anti-CD63 (primary), and anti-CD35 (secretory) or appropriate isotype controls in FACS media for 20 min on ice. Cells were washed following staining and resuspended in FACS media for analysis. Cells were gated for live cells (Live/Dead-negative) and then neutrophils (CD11b- and Ly6G-positive).

The median fluorescence intensity (MFI) was quantified for surface CD63 and CD35 relative to staining with an isotype control antibody.

#### Membrane Polarization Flow

After an hour rest, neutrophils were stained with 250 nM DiBAC4(3) for 8 minutes at 37°C then stimulated for ten minutes and run on the flow cytometer. The median fluorescence intensity (MFI) was quantified for DiBAC4(3) and normalized to unstimulated neutrophils.

### Confocal Microscopy

NET Microscopy Neutrophils were stained with CellMask deep red (5 mg/mL) and incubated at 37°C for 30 min. Cells were washed and 4×10^5^ neutrophils were plated in 150 uL of D10 on a poly-L Lysine coated 8 Well Chambered Cover Glass. After 1 hr incubation at 37°C, 5% CO2 cells were stimulated with Baclight Red-stained strains of *S. aureus* for indicated times. Hoescht 33342 (10 mM) was added during the last 30 min of incubation. Following stimulation the media was aspirated and neutrophils fixed with 4% PFA for 20 min ProLong Gold Antifade reagent was then applied. For HCQ (30pM) treatment, neutrophils were pretreated for 30 min prior to stimulation with *S. aureus*. NETs were imaged on a ZEISS LSM 900 with Airyscan. 2 confocal microscope. Images were analyzed using ImageJ software.

### Anti-dsDNA IgG ELISA

Vinyl 96-well plates were coated with dsDNA (10µg/mL) and incubated overnight at 4°C. The wells were washed before and after blocking and samples/standards were added before incubating overnight at 4°C. Captured antibodies were detected with anti-mouse IgG-alkaline phosphatase, A standard curve was generated using α-dsDNA and antibody. Absorbance at 405 nm for each well was quantified using a BioTek Cytation plate reader. Samples and standards were read in duplicate and DNA abundance was quantified relative to the standard.

### IFNα ELISA

In-vitro supernatant samples and serum samples were diluted 1:3 and 1:10 respectively in assay dilution buffer. Total IFNα protein was quantified using Mouse IFNα ProQuantum Immunoassay Kit following manufacturer instructions. Plates were ran on a Quantstudio 3 (Applied Biosystems).

### Mitochondrial Protein Immunoblot

Murine neutrophils were pooled from 2 mice and mitochondria were isolated from the cytosol (Thermo Scientific, 89874) per manufacturer instructions. Mitochondrial fractions were lysed using 1X RIPA containing protease inhibitors. Mitochondrial lysates were heated for 5 minutes at 100°C in the presence of Laemmli loading buffer containing β-mercaptoethanol. A volume of 25 µL/well was loaded onto a 15% polyacrylamide gel, followed by electrophoresis. Gels were transferred onto nitrocellulose using a Trans-Blot Turbo and blocked at 4°C with Intercept (TBS) blocking buffer for 4 h before an overnight incubation in either primary LDHB or SOD2 diluted at 1:1,000 in TBST with 1% milk. Secondary antibody incubations were done for 4 hours with 1:10,000 goat anti-rabbit IgG HRP before developing with a Chemiluminescent Substrate kit (Thermo Scientific, 34580) per manufacturer instructions. Blots were then imaged on an iBright 1500 and densitometry quantified by ImageJ software.

### Fluorescent Plate Reader Assays

#### Calcium

Murine neutrophils were isolated and resuspended in clear D10 and stained with Fluo-4 calcium dye (1:2,000) for 40 minutes and Hoechst dye (1:1000) for 20 minutes. The cells were washed and plated (250,000/well) in clear D10 media in a black-wall, clear bottom, 96-well plate that was Poly-L Lysine coated. After an hour rest at 37°C, 5% CO_2_, fluorescence intensity of total DNA (Hoescht 33342) and calcium were read on a Biotek Cytation plate reader. Neutrophils were stimulated with bacteria at an MOI of 10 and Hoechst 33342 and Fluo-4 fluorescence was read every 5 minutes for 1 hour.

#### Characterization of A23187 and PMA induced NETs

Murine neutrophils were isolated and plated (500,000/well) on a Poly-L Lysine coated, black wall, clear bottom plate. A23187 (5 uM) and PMA (50 nM) were added to the neutrophils and the plate was cultured for 16 hours at 37°C, 5% CO_2_. Wells were stained with SYTOX blue (500 nM) to quantify total DNA and antibodies specific to MPO, NE, and H3Cit with appropriate fluorescent secondary antibodies to quantify protein abundance.

### Dilution Spot Plate Assays

#### Neutrophil killing assay

Neutrophils were seeded at 2×10^5^ cells/well in a 96-well plate and were allowed to rest for 30 min in a 37°C, 5% CO_2_ incubator. Neutrophils were challenged with *S. aureus* opsonized in 1:1 DMEM:non-heat inactivated FBS at a MOI = 0.5. After 4 hours, samples were serially diluted and spot-plated onto TSA. Percent growth was quantified relative to bacteria grown in the absence of neutrophils.

#### NET killing assay

Murine neutrophils were isolated and plated (500,000/well) on a Poly-L Lysine coated, black wall, clear bottom plate. A23187 (5 uM) and PMA (50 nM) were added to the neutrophils and the plate was cultured for 16 hours at 37°C, 5% CO_2_. Neutrophils were challenged with *S. aureus* at a MOI = 0.2. After 4 hours, samples were serially diluted and spot-plated onto TSA. Percent growth was quantified relative to bacteria grown in the absence of neutrophils.

### Quantitative PCR

RNA was extracted from murine neutrophils using the Qiagen RNeasy mini kit following an enzymatic digestion with lysis buffer supplemented with lysozyme (2.88 mg/mL), mutanolysin (2.4 U/µL), proteinase K (3.85 mg/mL), and EDTA (19.2 mM). RNA was further purified by performing a second DNAse treatment with Turbo DNAse kit following the manufacturer’s protocol. RNA was quantified using a qubit RNA HS kit. 318 ng of RNA was then reverse transcribed to cDNA using the SuperScript III First-Strand Synthesis Kit. Quantitative PCR (qPCR) was performed using TaqMan assays to quantify transcript levels of LDHa (Mm01612132_g1 FAM), LDHb (Mm01267402_m1), and IFNα (Mm03030145_gH). Assays were prepared in triplicate with the TaqMan Gene-Expression Master Mix, and the plates were run on the QuantStudio 3 Real-Time PCR System (Applied Biosystems) following manufacturer instructions. Relative transcript abundance was calculated using the 2−ΔΔCt method, normalizing each target’s cycle threshold (Ct) to the endogenous control gene, ACTB (Mm02619580_g1 VIC).

### Statistics

Specific statistical details for each experiment can be found in the corresponding figure legend including statistical tests used, exact value of n, what n represents, definition of center, and dispersion and precision measures. Error bars for all experiments represent standard error of the mean. A minimum of three experimental replicates were performed for each assay and the specific number of replicates is noted in the corresponding figure and figure legend. Statistical work was performed using Prism 6 software (GraphPad) and significance is indicated on the graphs as follows: ^∗^, P ≤ 0.05; ^∗∗^, P ≤ 0.01; ^∗∗∗^, P ≤ 0.001; ^∗∗∗∗^, P ≤ 0.0001; ns, not significant.

## Acknowledgements

We thank Eric Skaar, Megan Kiedrowski, Carolyn Ibberson, and Alexander Horswill for providing bacterial strains. We thank Amit Joshi for providing the confocal microscopes. We also thank all the members of the Monteith and Sparer labs who provided feedback on the project and paper. We thank the individuals who provided blood and the support staff at UTMC who facilitated the process. We thank Trixie TenBarge and Onyx Mitchell for providing emotional support.

## Author Contributions

Conceptualization: E.G.T., J.D.B., and A.J.M. Methodology: E.G.T., A.D.W., M.L.H., and A.J.M. Investigation: E.G.T., A.D.W., M.L.H., H.A.H., H.E., B.E.H., E.B., E.C.M., S.R.M., N.M.V. C.C.L., J.M.W., and J.F. Data curation: E.G.T., A.D.W., M.L.H., J.M.W., J.F., and A.J.M. Resources: T.E.S., L.J.C., J.D.B., and A.J.M. Funding acquisition: A.J.M. Project administration: T.E.S., L.J.C., J.D.B., and A.J.M. Supervision: J.F., T.E.S., L.J.C., J.D.B., and A.J.M. Writing-original draft: E.G.T. and A.J.M. Writing-review and editing: E.G.T., A.D.W., M.L.H., H.A.H., H.E., B.E.H., E.B., E.C.M., S.R.M., N.M.V. C.C.L., J.M.W., J.F., T.E.S., L.J.C., J.D.B., and A.J.M.

## Materials availability

All data and unique reagents are available upon request to the corresponding author.

