## Supplemental Figures for "A TLR7/9-IFNα-LDHB axis drives vital NET release and compromises antibacterial defense in lupus"

**Supplemental Data**

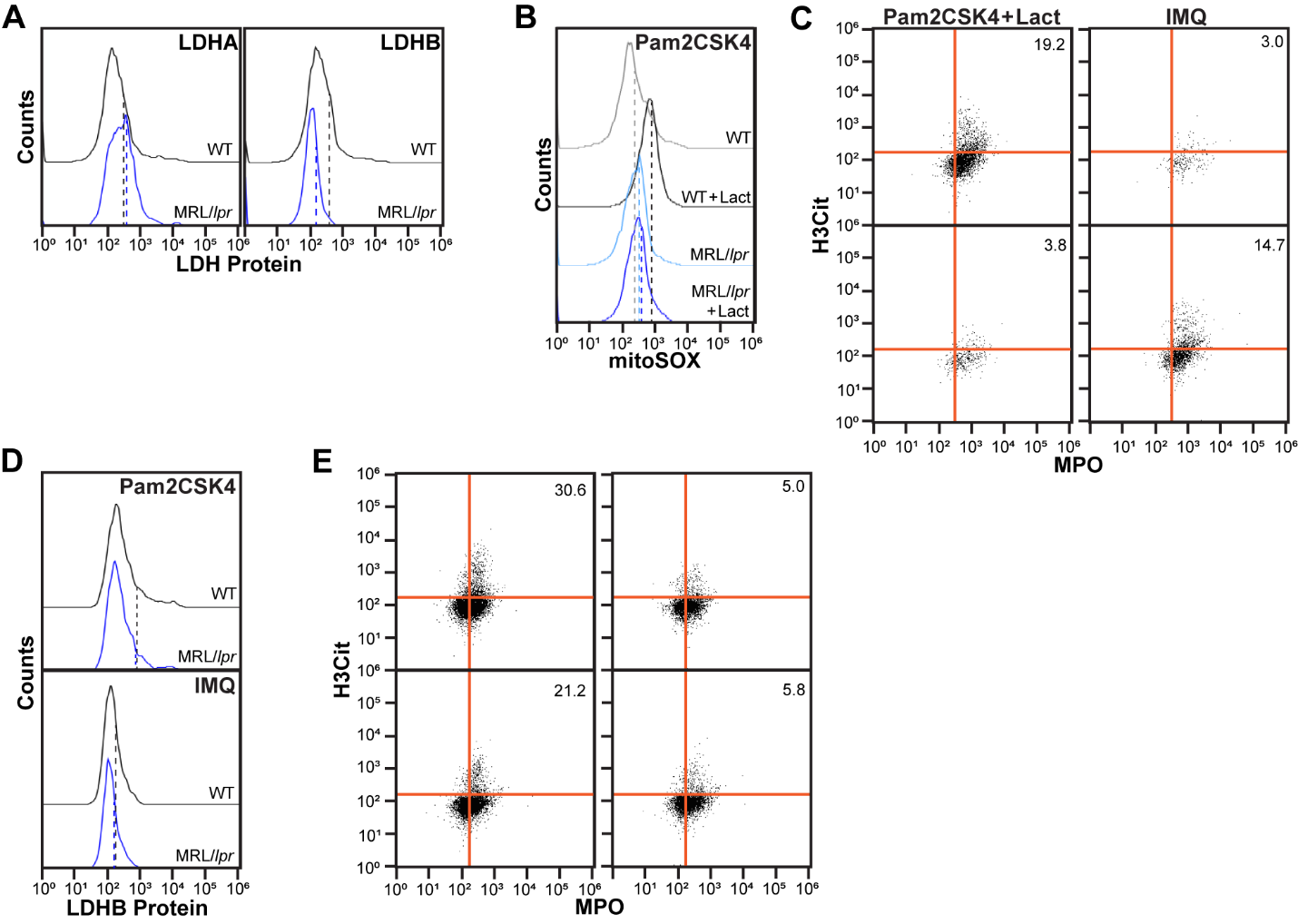

**Supplemental Figure 1: TLR7/9 signaling suppresses LDHB and impairs lactate-driven suicidal NETosis in lupus-prone neutrophils.** (**A**) Neutrophils were isolated from WT or MRL/*lpr* and total LDHA or LDHB protein were quantified by flow cytometry. (**B-C**) WT or MRL/*lpr* neutrophils were treated with differing TLR agonists in the presence or absence of lactate (1 mM) and (*B*) mitochondrial ROS or (*C*) suicidal NETosis were quantified by flow cytometry. (**D**-**E**) Neutrophils were stimulated with TLR agonist (50 ng/mL) and IFNγ (50 ng/mL) and cultured for 16 hours. Neutrophils were cultured with *S. aureus* for 3 hours and (**D**) total LDHB protein or (**E**) suicidal NETosis were quantified by flow cytometry. (*A-B, D*) Dashed line = MFI.

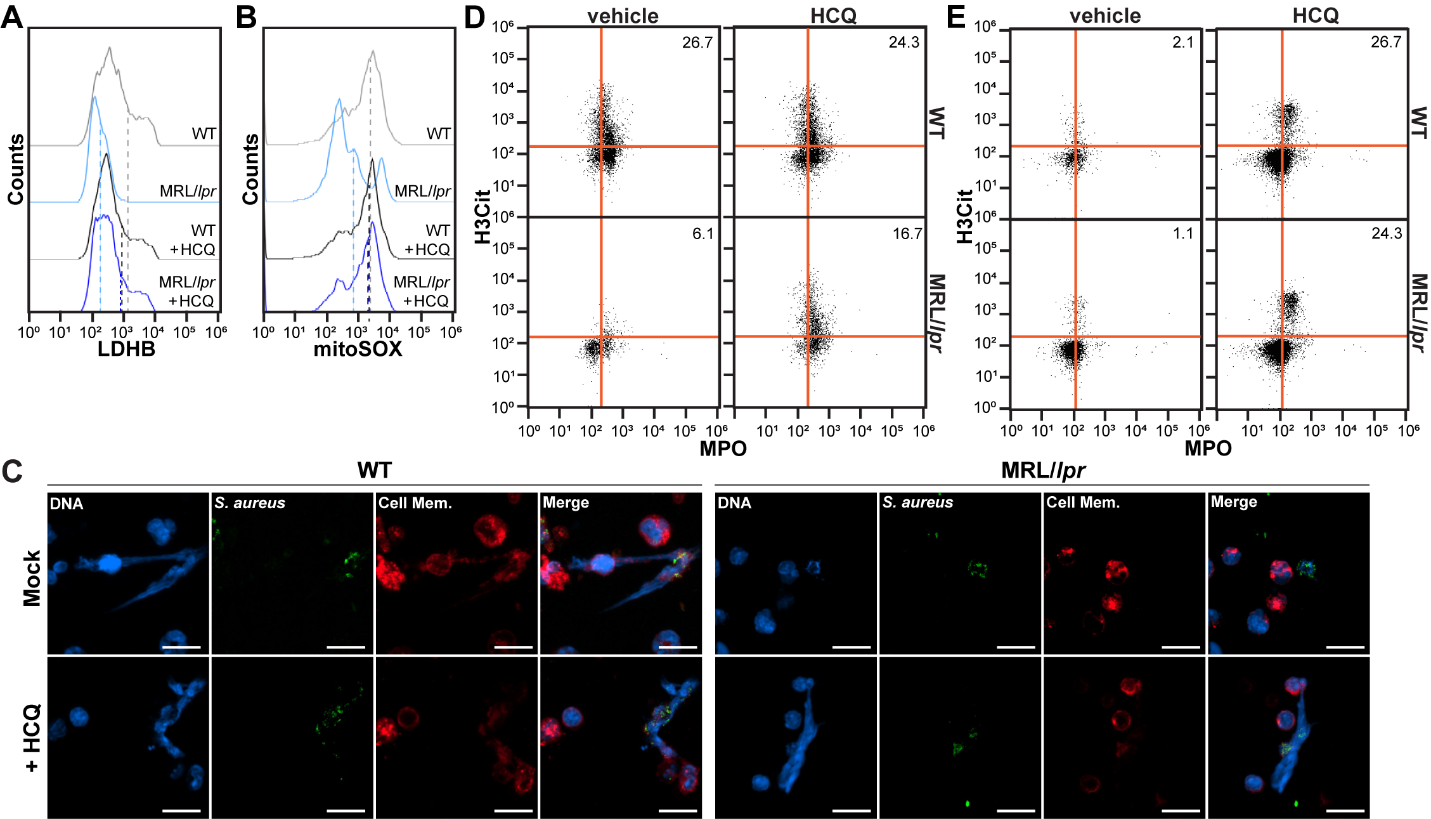

**Supplemental Figure 2: HCQ restores mitochondrial ROS and suicidal NETosis but fails to prevent chronic vital NET release in MRL/*lpr* neutrophils.** WT or MRL/*lpr* mice were mock treated or treated with HCQ (40 mg/kg) daily for 2 days. Neutrophils were isolated from the bone marrow, cultured with *S. aureus* for 3 hours, and (*A*) total LDHB protein, (*B*) mitochondrial ROS, neutrophils undergoing (*D*) suicidal NETosis or (*E*) vital NET release were quantified by flow cytometry. (*C*) Deposition of NETs were also quantified by confocal microscopy (bar = 10 µm). (*A-B*) Dashed line = MFI.

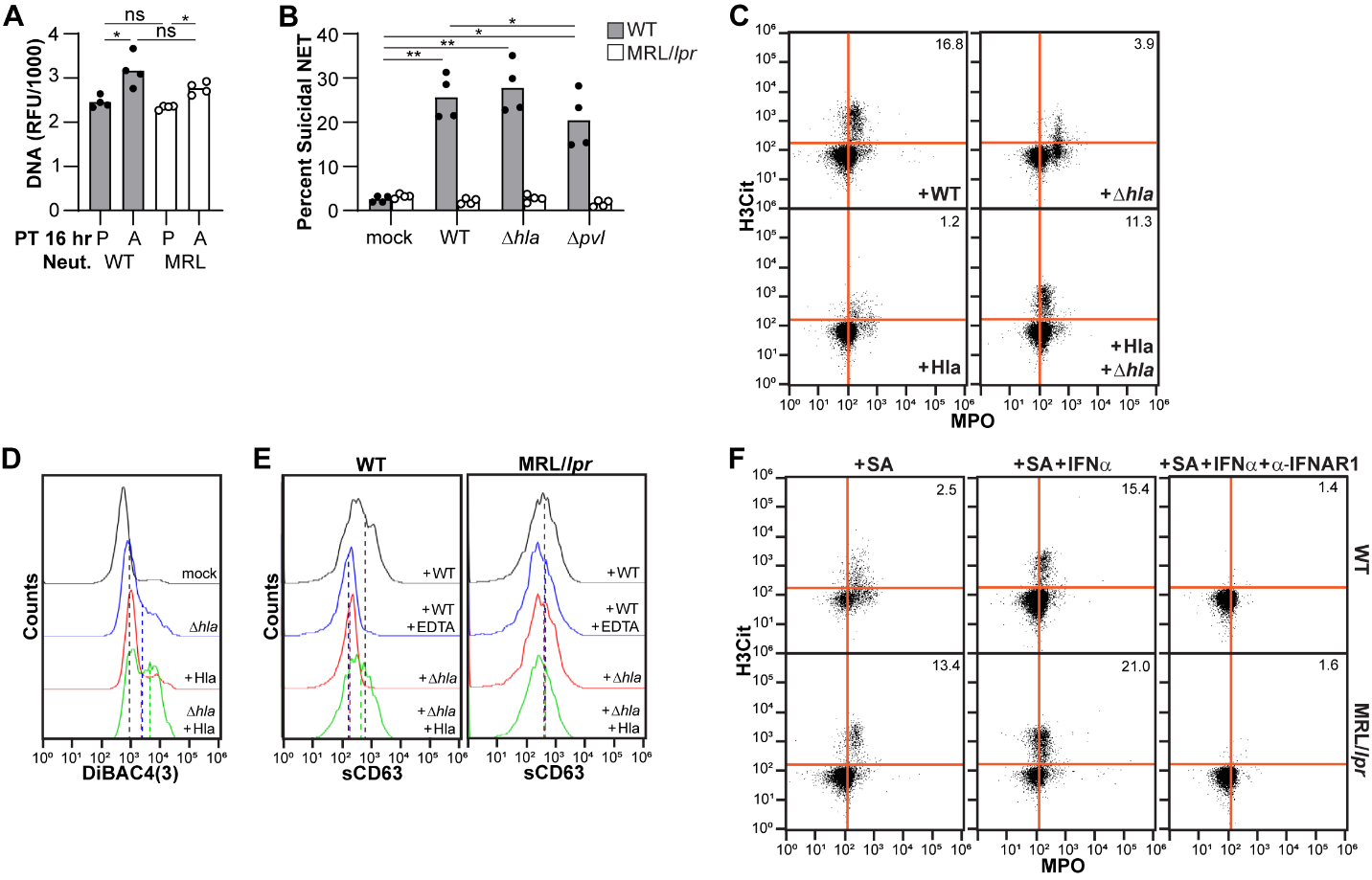

**Supplemental Figure 3: Hla expression does not effect suicidal NETosis.** (**A**) WT and MRL/*lpr* neutrophils were treated with PMA or A23187 for 16 hours and the resulting NETs subsequently stained to quantify DNA. (**B**) WT or MRL/*lpr* neutrophils were cultured with WT, Δ*hla*, Δ*pvl,* or *Δhla* + *hla* *S. aureus* in the presence or absence of Hla protein and suicidal NETosis was quantified after 3 hours by flow cytometry. (**C**) WT neutrophils were cultured with WT or Δ*hla* *S. aureus* in the presence or absence of Hla protein and vital NET release was quantified after 30 minutes by flow cytometry. (**D**) WT neutrophils were loaded with DiBAC_4_(3) and stimulated with Δ*hla* and/or Hla protein. The median fluorescence intensity (MFI) was quantified by flow cytometry and normalized to unstimulated neutrophils. (**E**) WT or MRL/*lpr* neutrophils were cultured with WT or Δ*hla* *S. aureus* in the presence or absence of EDTA or Hla protein. After 30 minutes, neutrophils were stained for surface expression of CD63 (primary granules) and quantified by flow cytometry. (**F**) WT or MRL/*lpr* neutrophils were treated with IFNα, and/or an α-IFNAR1 antibody and cultured with *S. aureus*. After 3 hours, vital NET release was quantified by flow cytometry. (*A-B*) Each point represents neutrophils isolated from a single mouse. (*A*) One- or (*B*) two-way ANOVA with Tukey multiple comparisons test (**p* ≤ 0.05, ***p* ≤ 0.01, ns = not significant). (*D-E*) Dashed line = MFI.

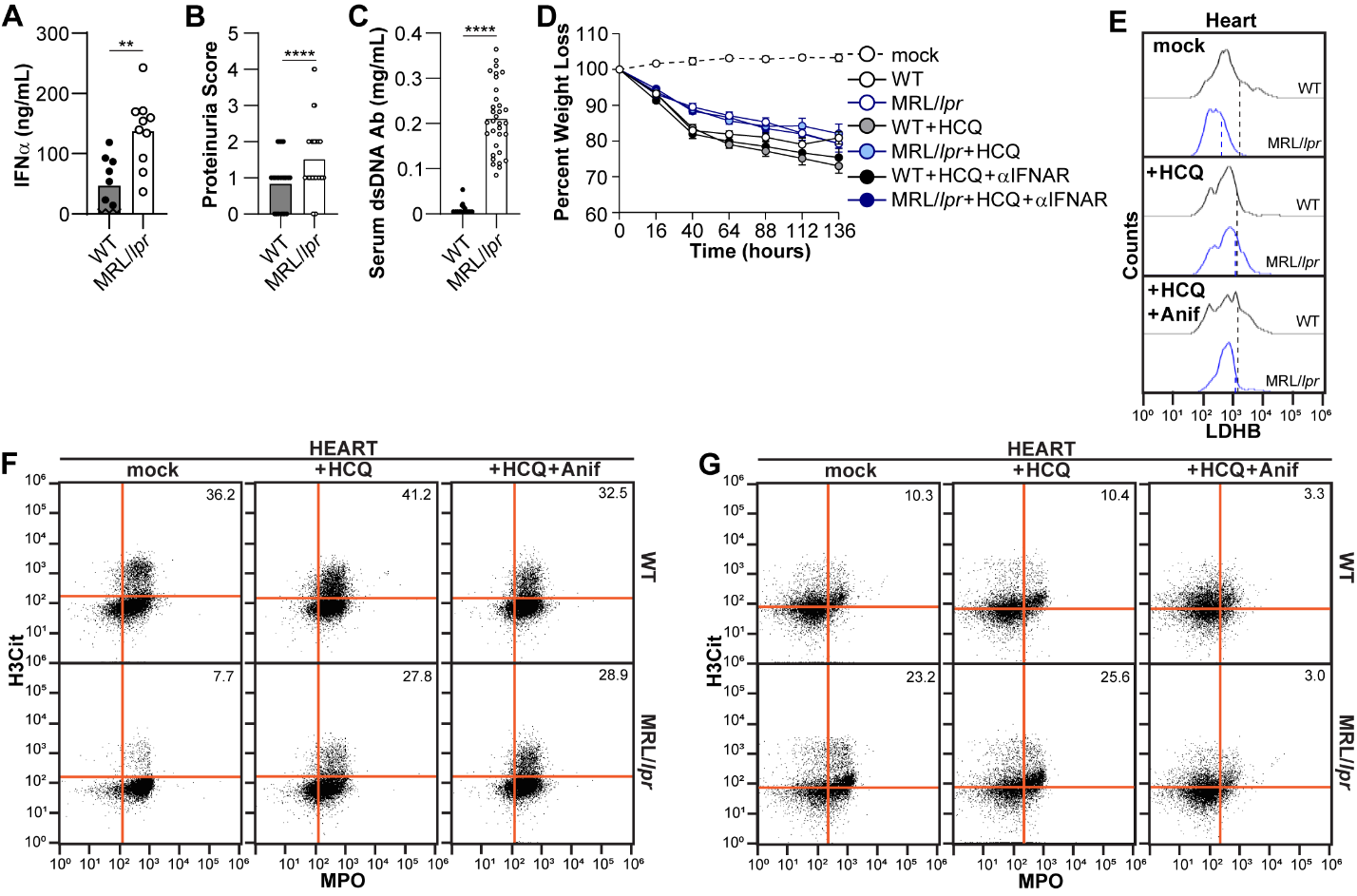

**Supplemental Figure 4: MRL/*lpr* have increased IFNα during infection.** WT or MRL/*lpr* mice were retro-orbitally infected with *S. aureus* and (**A**) 6 days post-infection, serum was isolated and IFNα quantified by ELISA. (**B**-**C**) The day before infection, (*B*) proteinuria scores were assigned by quantifying protein levels in the urine (0=Trace, 1=30 mg/dL, 2=100 mg/dL, 3=300 mg/dL, 4=2000+ mg/dL) and (*C*) IgG levels were quantified from the serum of the mice by ELISA. (**D**) Weight loss was monitored during infection. Percent weight loss is normalized to starting weight. Neutrophils from the heart were analyzed for (**E**) LDHB protein expression, (**F**) suicidal NETosis, and (**G**) vital NET release by flow cytometry. (*A*-*C*) Each point represents a single mouse or (*D*) the mean of all mice. (*A-C*) Unpaired t test or (*D*) error bars reflect standard error (**p ≤ 0.01,****p ≤ 0.0001). (*E-F*) Dashed line = MFI.

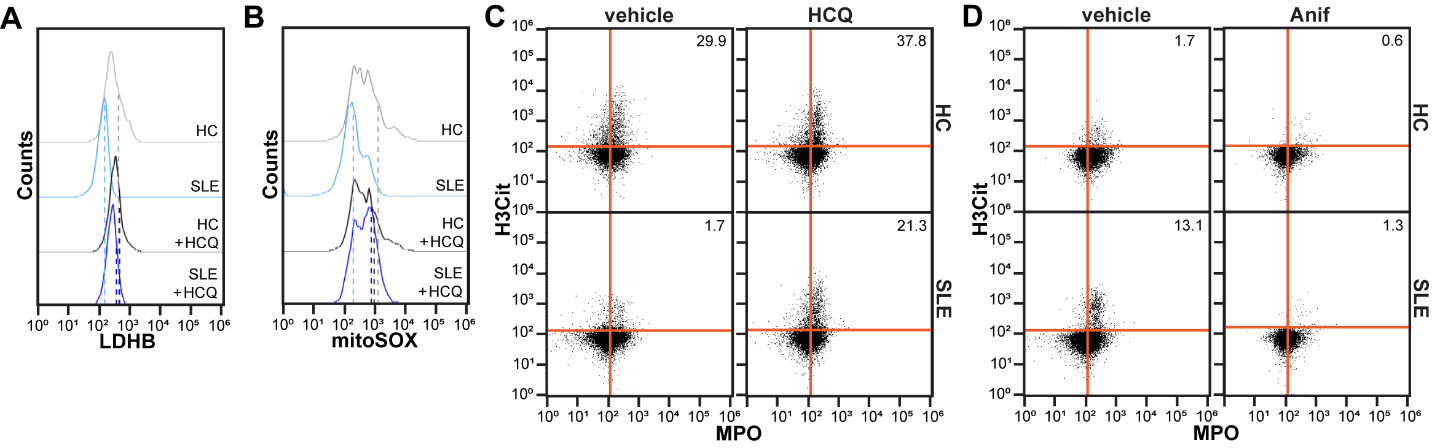

**Supplemental Figure 5: HCQ and anifrolumab restore NET subtype regulation in SLE neutrophils.** Neutrophils were isolated from HC or SLE patients. (**A**) LDHB protein expression in unstimulated neutrophils or, (**B**) mitochondrial ROS, (**C**) suicidal NETosis, (**D**) vital NET release in response to *S. aureus* were quantified by flow cytometry. (*A-B*) Dashed line = MFI.

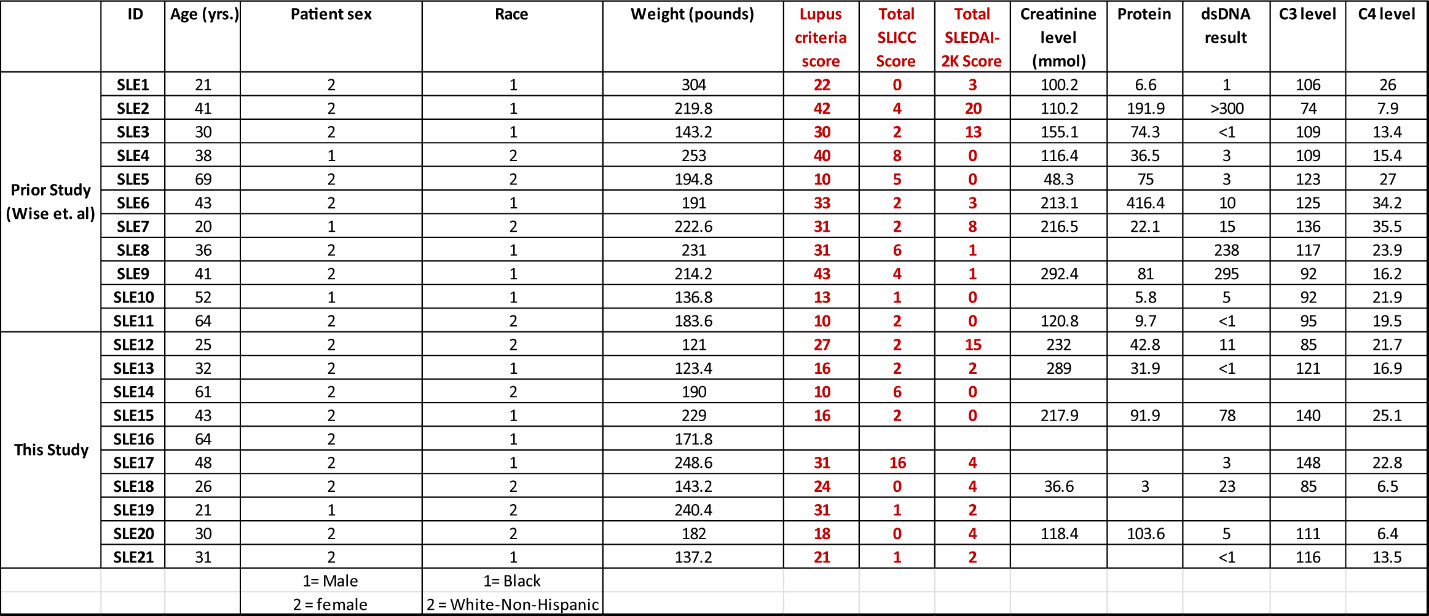

**Supplemental Table 1: SLE donor demographics and disease state.**

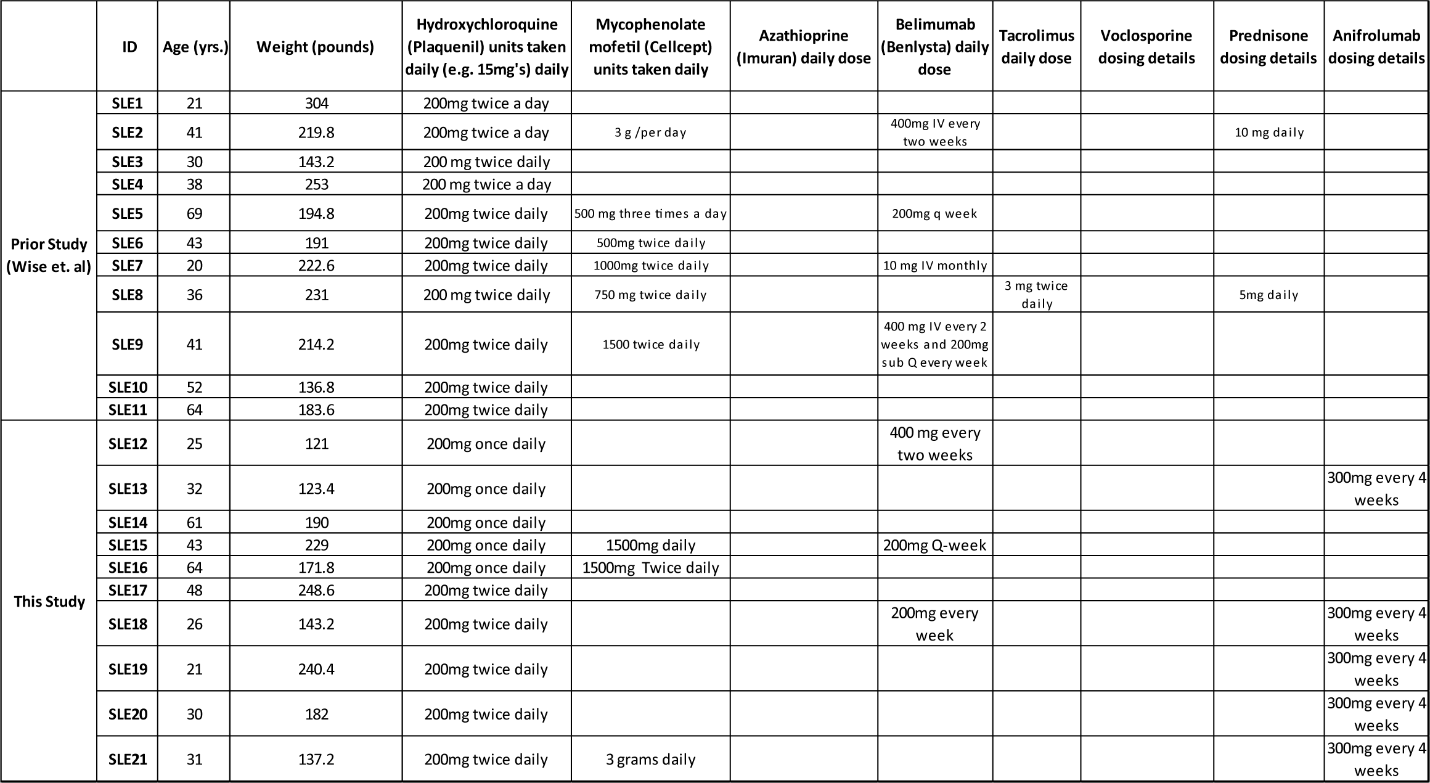

**Supplemental Table 2: Medications taken by the SLE donors.**
